# Mitochondrial ROS and HIF-1α signalling mediate critical period plasticity

**DOI:** 10.1101/2024.11.22.624842

**Authors:** Daniel Sobrido-Cameán, Bramwell Coulson, Michael Miller, Matthew C.W. Oswald, Tom Pettini, Richard A. Baines, Matthias Landgraf

## Abstract

As developing networks transition from spontaneous irregular to patterned activity, they undergo plastic tuning phases, termed “critical periods”; “critical” because disturbances during these phases can lead to lasting changes in network development and output. Critical periods are common to developing nervous systems, with analogous features shared from insects to mammals, yet the core signalling mechanisms that underly cellular critical period plasticity have remained elusive. To identify these, we exploited the *Drosophila* larval locomotor network as an advantageous model system. It has a defined critical period and offers unparalleled access to identified network elements, including the neuromuscular junction as a model synapse. We find that manipulations of a single motoneuron or muscle cell during the critical period lead to predictable, and permanent, cell-specific changes. This demonstrates that critical period adjustments occur at a single cell level. Mechanistically, we identified mitochondrial reactive oxygen species (ROS) as causative. Specifically, we show that ROS produced by complex I of the mitochondrial electron transport chain, generated by the reverse flow of electrons, are necessary and instructive for critical period-regulated plasticity. Downstream of ROS, we identified the *Drosophila* homologue of hypoxia inducible factor (HIF-1α), as required for transducing the mitochondrial ROS signal to the nucleus. This signalling axis is also sufficient to cell autonomously specify changes in neuronal properties and animal behaviour but, again, only when activated during the embryonic critical period. Thus, we have identified specific mitochondrial ROS and HIF-1α as primary signals that mediate critical period plasticity.

## Introduction

The emergence of network function is arguably among the least well understood aspects of nervous system development. It is well known that the emergence of network function, which occurs during late stages of nervous system development, correlates with a phase of adjustment during which networks transition from seemingly irregular to more patterned activity. These phases have been termed ‘sensitive’ or ‘critical periods’ and are commonly viewed as periods of heightened plasticity. Importantly, they are also highly sensitive to perturbations: errors induced during the critical period generally have lasting impact, with subsequent plasticity mechanisms unable to correct them^1–3^. Thus, “decisions” made during the critical period can specify neuronal properties over the long-term, and abnormal critical period experiences likely contribute to epilepsy and other neurodevelopmental conditions such as schizophrenia and autism spectrum disorder^4,5^. Though critical periods have been studied for decades, notably in mammalian sensory systems^3,6,7^, many questions remain. Here, we focused on two fundamental unresolved questions: whether critical periods are principally a network phenomenon, or a property of individual cells and, secondly, the identity of the primary signals directing critical period plasticity.

A difficulty in understanding critical periods has been the complexity of mammalian circuitry, including the realisation that different cell types can behave differently^6^. Therefore, we exploit a much simpler experimental system, the fruit fly, *Drosophila melanogaster*, which has enabled the identification of numerous genes and mechanisms that orchestrate the development and function of the nervous system^8–10^. Importantly, the larval locomotor network has a well-defined critical period in late embryogenesis, with features that suggest it is analogous to mammalian critical periods^11,12^. As is the case for mammals, *Drosophila* critical periods, both in the embryo and during pupal metamorphosis, are developmental windows that are fundamentally activity-regulated, with an important role for shaping the network inhibition:excitation balance^11–14^. For example in *Drosophila,* transient correction of network inhibition:excitation balance during the embryonic critical period effects a sustained rescue of seizure phenotypes otherwise caused by mutation of the single sodium channel^11,12^. Experimentally, this model system allows reliable access to identified cells of known connectivity and function. Unlike mammals, *Drosophila melanogaster,* as an insect, is not warm blooded, but poikilothermic, and is therefore vulnerable to external temperature variation. For the larval locomotor network, we found that increases in ambient temperature elevate network activity and, when applied during the embryonic critical period, phenocopy pharmacological and genetic activity manipulations. Thus, heat stress of 32°C, which is ecologically relevant to temperatures that *Drosophila* experience in the wild^15^, represents an ecologically relevant stimulus with which to study critical period biology in this system.

Temperature affects biochemical reactions in general, and metabolic and mitochondrial processes in particular^16^. In *Drosophila,* heat stress causes a reversal of the flow of electrons within the mitochondrial electron transport chain (ETC), from Complex-II back to Complex-I. This so-called reverse electron transport (RET) leads to the production of reactive oxygen species (ROS) at Complex-I, due to a partial reduction of oxygen and the formation of superoxide anions. In *Drosophila*, as well as mouse, RET-generated ROS have been linked to improved stress resistance and lifespan^17–21^. Here, we identified RET-based generation of ROS at mitochondrial Complex-I as a primary critical period signal that is necessary and sufficient for instructing changes in subsequent network development.

Downstream of mitochondrial ROS, we further identified the conserved Hypoxia-inducible factor 1 alpha (HIF-1α), of which *Drosophila* has only a single homologue, called *similar (sima)*^22–24^. HIF-1α is a sensor for both oxygen and ROS levels, and is a regulator of metabolism^25^. During normal development, HIF-1α regulates many processes, including neurogenesis, angiogenesis, and cell survival^26–28^ and its dysregulation is linked to a range of cancers^29^. In summary, we report that critical period plasticity is principally enacted cell autonomously; with metabolic mitochondrial ROS as a primary signal that is transduced by the conserved HIF-1α pathway, instructing subsequent development of cellular excitable and synaptic properties, and thus network stability and behavioural output.

## Results

### Mitochondrial ROS are plasticity signals during the embryonic critical period

The primary signals and pathways that underlie critical period plasticity remain elusive. To identify these, we used the *Drosophila* larval neuromuscular junction (NMJ) as a proven experimental model^8–10^ and, as a means to induce a robust phenotype, we used transient exposure to 32°C heat stress during a defined embryonic critical period (occurring at 17-19 hrs after egg laying at 25°C). We have previously shown this manipulation to increase network excitation (i.e. increased action potential firing) phenocopying the effects of direct activity manipulations^30^. Specifically, embryos experiencing 32°C heat stress during their critical period show subsequent change to larval development of the NMJ. This change is seemingly permanent and manifests as presynaptic terminal overgrowth (Figure 1A-C) and a decrease in the postsynaptic high-conductance glutamate receptor subunit, GluRIIA (Figure 1D), but not the lower conductance subunit, GluRIIB (Figure 1E).

**Figure 1.**
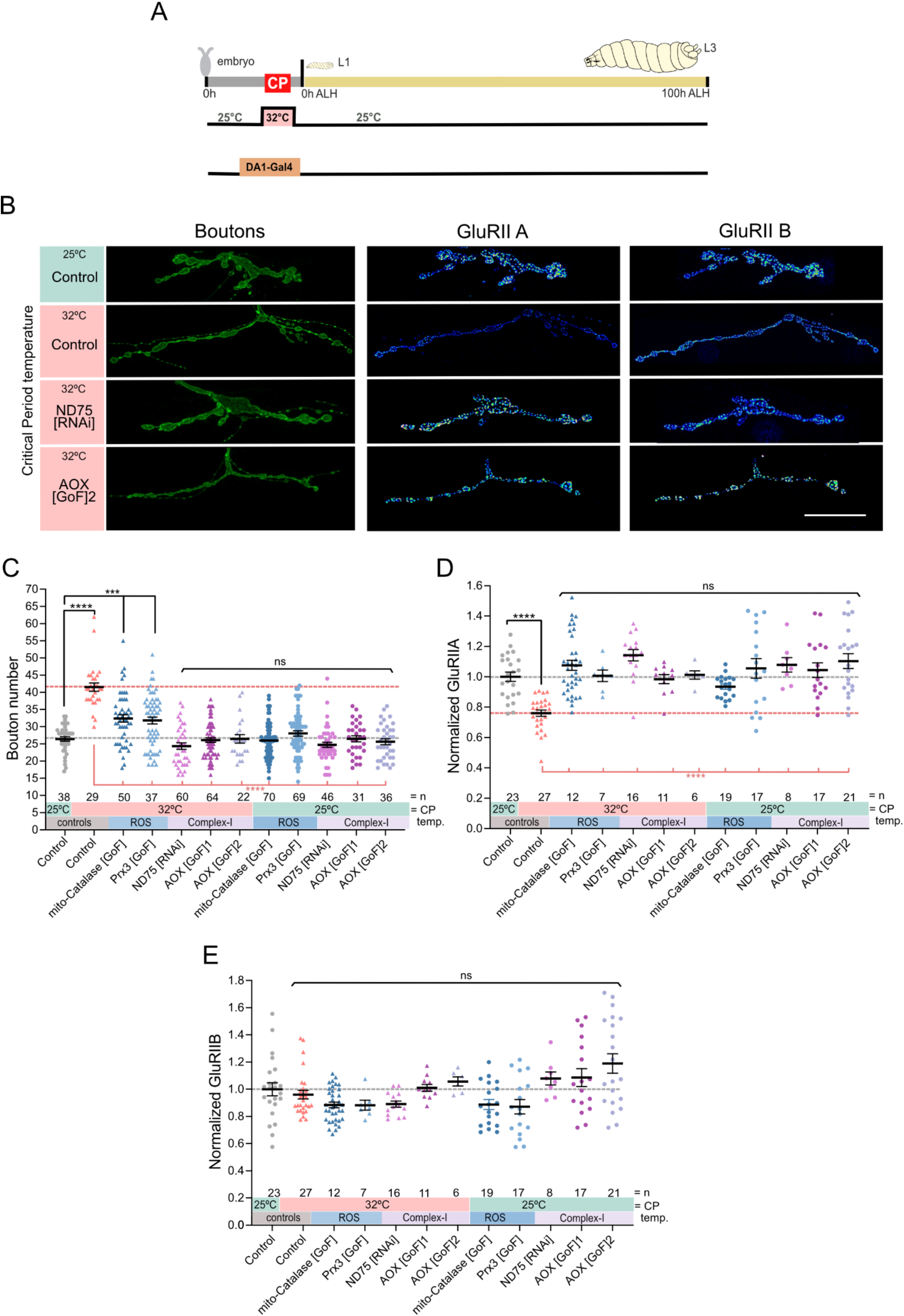
Mitochondrial ROS generated by reverse electron transport in muscles are necessary for critical period heat stress to change NMJ development. **A)** Experimental paradigm **B)** Heat stress experienced during the embryonic critical period (32°C *vs* 25°C control) leads to increased aCC NMJ terminal size and decreased postsynaptic GluRIIA, while not affecting subunit GluRIIB expression. Simultaneous genetic manipulation of muscle DA1 during embryonic stages only identifies mitochondrial ROS generated by reverse electron transport as necessary signals. “Control” indicates control genotype heterozygous for Oregon-R and DA1-Gal4. Larvae were reared at the control temperature of 25°C until the late wandering stage, 100 hrs after larval hatching (ALH). GluRIIA and GluRIIB subunits are displayed with lookup table “fire” to illustrate signal intensities (warmer colours indicating greater signal intensities). Scale bar = 20 µm. **C)** Dot-plot quantification shows changes to aCC NMJ growth on its target muscle DA1, based on the standard measure of the number of boutons (swellings containing multiple presynaptic release sites/active zones). Data are shown with mean ± SEM, ANOVA, *** p<0.0001, **** p<0.00001, ‘ns’ indicates statistical non-significance. Black asterisks indicate comparison with the control condition of 25°C throughout, genetically unmanipulated. Red asterisks indicate comparisons with control genotype exposed to 32°C heat stress during the embryonic critical period. **D)** Dot-plot quantification shows changes in levels of the GluRIIA receptor subunit at aCC NMJs quantified in C). Data are shown with mean ± SEM, ANOVA, **** p<0.00001, ‘ns’ indicates statistical non-significance. Black asterisks indicate comparison with the control condition of 25°C throughout, genetically unmanipulated. Red asterisks indicate comparisons with control genotype exposed to 32°C heat stress during the embryonic critical period. **E)** Dot-plot quantification as in D), but for the low conductance GluRIIB receptor subunit, which remains unaffected by these manipulations.

The NMJ overgrowth phenotype mirrors the outcome following neuronal overactivation-induced by reactive oxygen species (ROS) signalling, which we have previously studied^30,31^. We therefore asked if mitochondrial ROS might be involved. To test this, we genetically manipulated mitochondrial ROS levels in embryonic stages, selectively within a single muscle per half segment: the most dorsal muscle, dorsal acute 1 (DA1). Mis-expression of the mitochondria-targeted ROS scavengers, mito-Catalase or Perexiredoxin-3 (Prx3), in muscle DA1, cell-selectively rescues NMJ overgrowth (Figure 1C) and GluRIIA phenotypes that otherwise result from a 32°C heat stress during the embryonic critical period (Figure 1D). At the control temperature of 25°C, these same genetic manipulations have no such effects (Figure 1).

Mitochondria can generate ROS through various means, notably at Complex-I and Complex-III of the electron transport chain. Heat stress causes ROS generation at Complex-I in adult *Drosophila*, via reverse flow of electrons from Complex-II back to Complex-I, (termed ‘reverse electron transport’, or ‘RET’^19^). To test if 32°C heat stress in embryos causes RET-generated ROS in mitochondria, we cell-selectively knocked down a subunit of Complex-I, ND-75, required for RET-generated ROS. To complement this, we further neutralised RET-generated ROS by mis-expression of alternative oxidase (AOX), an enzyme found in the inner mitochondrial membrane of many organisms, though not vertebrates or insects^32–35^. In mitochondria, AOX acts as a bypass that transfers electrons from ubiquinone to oxygen, independent of Complexes-I and -III, thereby reducing the probability of electron backflow and subsequent ROS production by RET^19,36–38^. When we targeted either of these genetic manipulations to muscle DA1 during embryogenesis, both rescued the critical period heat stress phenotype (Figure 1). Under control conditions of 25°C both manipulations were neutral, leaving NMJ development unaffected (Figure 1). Together, our results show that during the embryonic critical period, RET-generated ROS at Complex-I are necessary for heat stress to effect a lasting change to NMJ development.

We next asked if ROS generated by Complex-I might also be sufficient to induce critical period plasticity in absence of systemic heat stress. To test this, we cell-selectively induced RET-ROS at Complex-I by mis-expression of yeast NADH dehydrogenase 1 (Ndi1). Ndi1 transfers electrons from NADH to the CoQ pool. This can lead to the accumulation of reduced CoQ and transfer of electrons to Complex-I, causing RET and associated ROS generation^28^. Mis-expression of Ndi1 in *Drosophila* has previously been shown to increase ROS production in a way indistinguishable from RET^20^. We found that transient embryonic mis-expression of Ndi1 in muscle DA1, at the control temperature of 25°C, cell-selectively phenocopies the characteristic critical period heat stress phenotype of NMJ overgrowth (Figure 2A-C) and reduced GluRIIA levels (Figure 2E). Thus, our results show that RET-generated ROS, generated from mitochondria during the embryonic critical period, are both necessary and sufficient for inducing lasting change to NMJ development.

**Figure 2:**
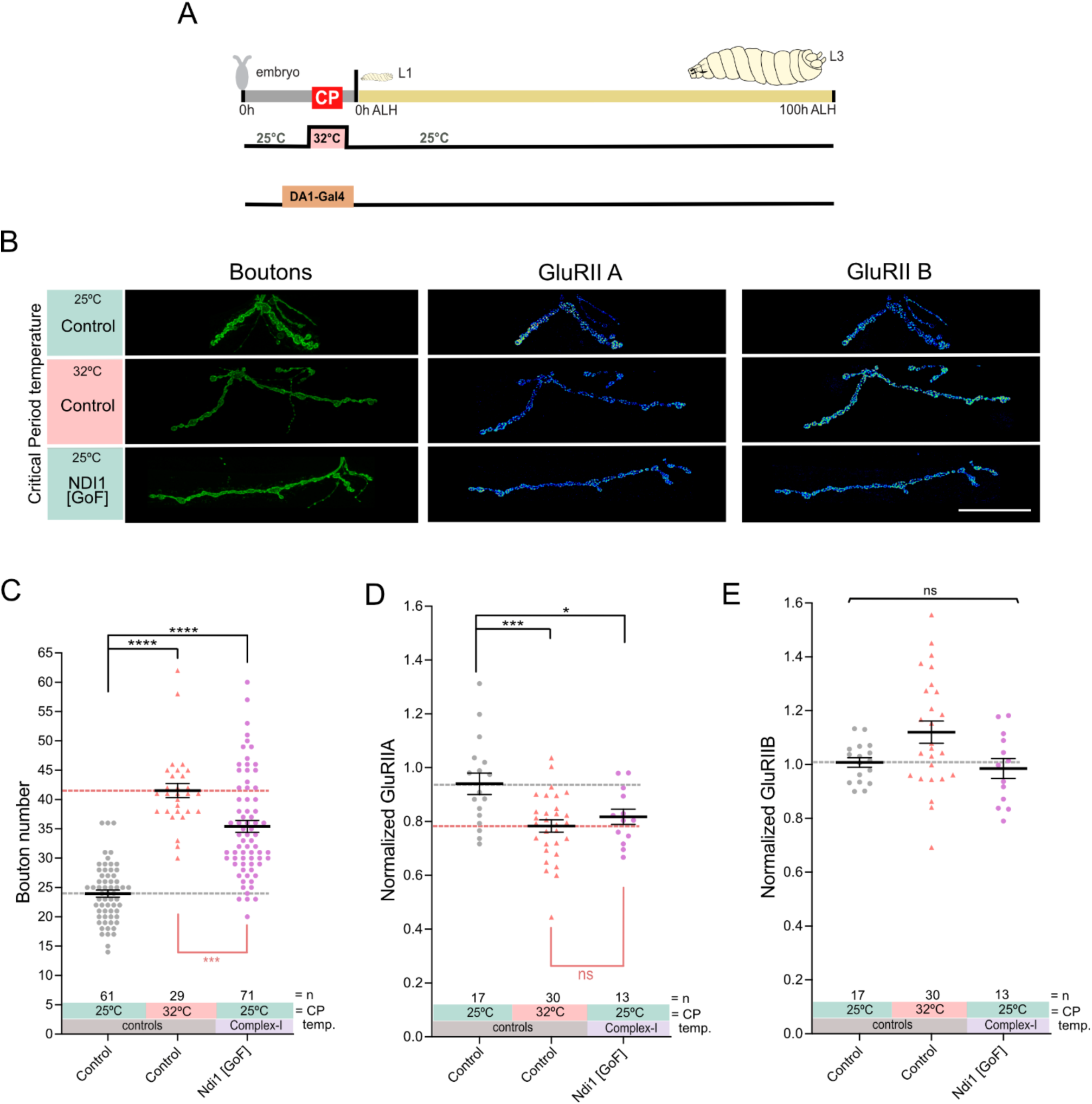
ROS by Complex-I, during the critical period, are sufficient to induce lasting changes to NMJ development. **A)** Experimental paradigm **B)** Induction of ROS at Complex-I via transient mis-expression of NDI1 in muscle DA1 during embryonic stages only phenocopies the effects that critical period heat stress has on subsequent NMJ development. “Control” indicates control genotype heterozygous for Oregon-R and DA1-Gal4. Larvae were reared at the control temperature of 25°C until the late wandering stage, 100 hrs ALH. GluRIIA and GluRIIB subunits are displayed with lookup table “fire” to illustrate signal intensities (warmer colours indicating greater signal intensities). Scale bar = 20 µm. **C)** Dot-plot quantification shows changes to aCC NMJ growth on its target muscle DA1, based on the standard measure of the number of boutons (swellings containing multiple presynaptic release sites/active zones). Data are shown with mean ± SEM, ANOVA, *** p<0.0001, **** p<0.00001. Black asterisks indicate comparison with the control condition of 25°C throughout, genetically unmanipulated. Red asterisks indicate comparisons with control genotype exposed to 32°C heat stress during the embryonic critical period. **D)** Dot-plot quantification shows changes in levels of the GluRIIA receptor subunit at aCC NMJs quantified in C). Data are shown with mean ± SEM, ANOVA, ** p<0.001, *** p<0.0001, ‘ns’ indicates statistical non-significance. Black asterisks indicate comparison with the control condition of 25°C throughout, genetically unmanipulated. Red asterisks indicate comparisons with control genotype exposed to 32°C heat stress during the embryonic critical period. **E)** Dot-plot quantification as in D), but for the low conductance GluRIIB receptor subunit, which remains unaffected by these manipulations.

### Mitochondrial RET-generated ROS are critical period signals for NMJ development

Synapse development requires close interactions between pre- and postsynaptic partners. Therefore, we next tested the roles of mitochondrial ROS in a presynaptic motoneuron, termed ‘aCC’, whose NMJ forms on muscle DA1. We identified a transgenic line that transiently targets Gal4 expression to the aCC presynaptic motoneuron selectively during embryonic stages, including the critical period. Using this “aCC-Gal4[1]” genetic tool (aka *RN2-O-Gal4*), we found that mis-expression of AOX in aCC motoneurons, to neutralise RET-induced ROS generation at Complex-I, was sufficient to rescue the NMJ overgrowth that normally results from embryonic 32°C heat stress (Figure 3 A-C). Critical period-induced increases in NMJ size led to a proportionate increase in the number of presynaptic active zones, the sites of vesicular neurotransmitter release (Figure 3D). However, simultaneous embryonic mis-expression of AOX in the aCC motoneurons fully rescues active zone number (Figure 3D). Conversely, transient embryonic generation of RET-ROS in the aCC motoneurons, by mis-expression of Ndi1, was sufficient to cell-selectively mimic the critical period heat stress phenotype (Figure 3B-D). These findings demonstrate that rescue could occur through either presynaptic or postsynaptic processes.

**Figure 3:**
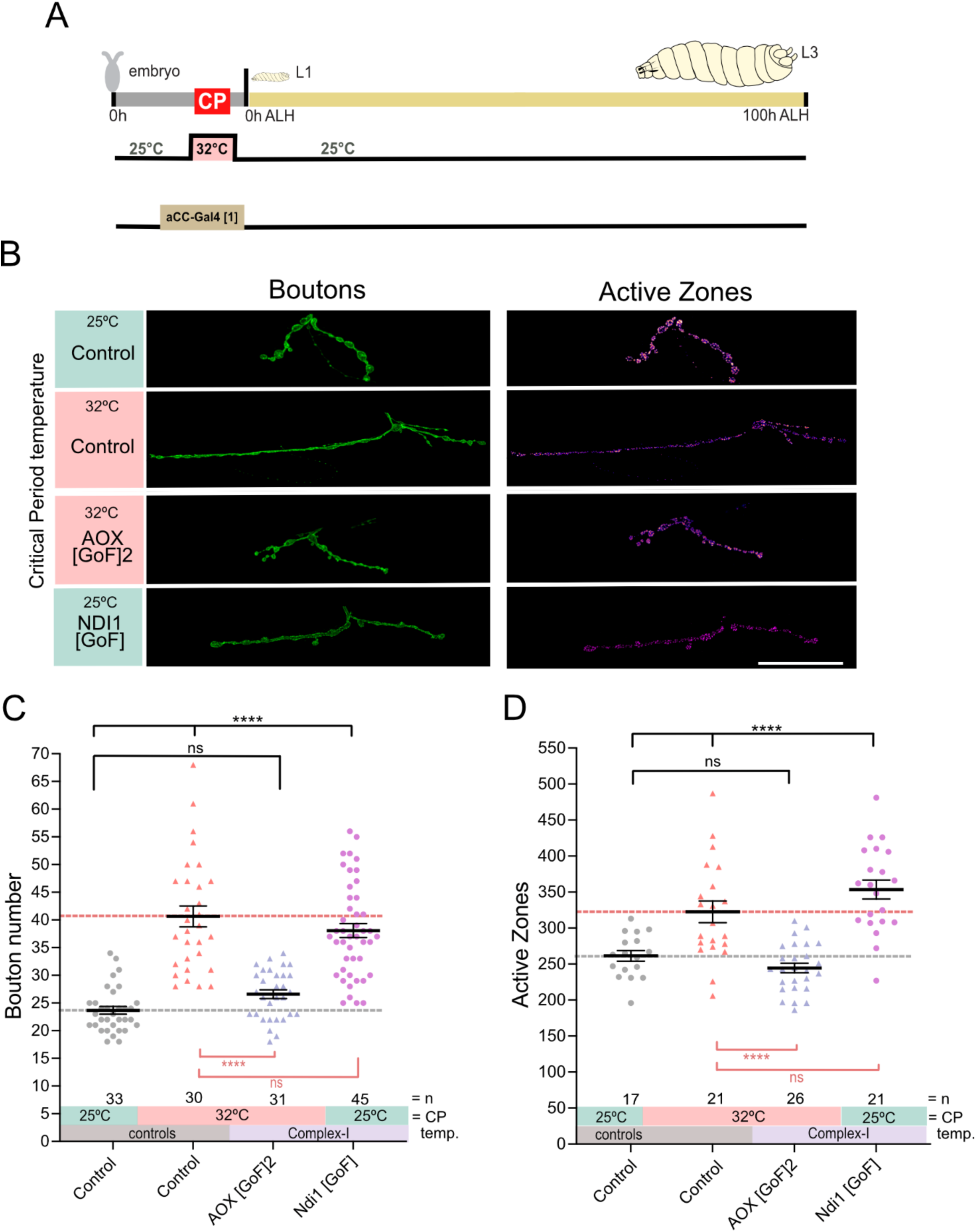
ROS by RET in embryonic motoneurons instructs subsequent NMJ development. **A)** Experimental paradigm **B)** Temperature experienced during the embryonic critical period (25°C control *vs* 32°C heat stress) and simultaneous, transient genetic manipulation of motoneuron aCC during embryonic stages only via aCC-Gal4 [1] (aka*RN2-O-Gal4*). “Control” indicates control genotype heterozygous for Oregon-R and aCC-Gal4[1]. Larvae were reared at the control temperature of 25°C until the late wandering stage, 100 hrs ALH. Scale bar = 20 µm. **C)** Dot-plot quantification shows changes to aCC NMJ growth on its target muscle DA1, based on the standard measure of the number of boutons (swellings containing multiple presynaptic release sites/active zones). Data are shown with mean ± SEM, ANOVA, **** p<0.00001, ‘ns’ indicates statistical non-significance. Black asterisks indicate comparison with the control condition of 25°C throughout, genetically unmanipulated. Red asterisks indicate comparisons with control genotype exposed to 32°C heat stress during the embryonic critical period. **D)** Dot-plot quantification shows changes in number of the active zones at aCC NMJs quantified in C). Data are shown with mean ± SEM, ANOVA, **** p<0.00001, ‘ns’ indicates statistical non-significance. Black asterisks indicate comparison with the control condition of 25°C throughout, genetically unmanipulated. Red asterisks indicate comparisons with control genotype exposed to 32°C heat stress during the embryonic critical period.

Next, we tested the importance of developmental timing during which transient RET-generated ROS can cause such lasting change, i.e. whether mitochondrial ROS are indeed critical period-associated signals. To this end, we contrasted the effects of transient genetic manipulations of aCC motoneurons, during the late embryonic stage, including the critical period, *versus* identical manipulations after the critical period has closed (i.e. during the first larval stage). We achieved temporal and spatial control of Gal4 expression by combining two transgenes: another, independently generated expression line, “aCC-Gal4[2]” (aka *GMR96G06-Gal4*), which specifically targets Gal4 to the aCC motoneuron during embryonic and also larval stages^39,40^, combined with a conditional Gal4 inhibitor from the recently published “AGES” system^41^. To induce Gal4 activity during the critical period, we fed auxin to gravid females. While for later expression, post critical period closure, we fed auxin to freshly hatched larvae for 24 hours. The results confirm that the timing of mitochondrial RET-generated ROS is important: only when cells generate mitochondrial ROS signals during the embryonic phase, including the critical period, does ROS cause lasting change, in keeping with this being a critical period of locomotor network development (supplementary figure 1).

### The conserved HIF-1α pathway transduces the mitochondrial ROS signal to the nucleus

We next considered how mitochondrial ROS, triggered by heat stress, might effect change to developmental outcomes. HIF-1α is a transcription factor that plays a central role in cellular responses to hypoxia and other stresses, including oxidative stress^42^. First, we knocked down the single *Drosophila* homologue of *HIF-1α*, *sima,* selectively in DA1 muscles while exposing embryos to 32°C during the critical period. This showed that HIF-1α is necessary to cause heat stress-induced critical period plasticity: NMJ overgrowth (Figure 4A-C) and a postsynaptic decrease of the GluRIIA receptor subunit (Figure 4D). Conversely, transient embryonic mis-expression of *HIF-1α* in DA1 muscles, at the control temperature of 25°C, is sufficient to induce these NMJ phenotypes (Figure 4). Under normoxic conditions, HIF-1α is continuously degraded by the proteasome, regulated via hydroxylation by the conserved prolyl hydroxylase, PHD (a single *Drosophila* homologue, *Hph/fatiga*). Hydroxylated HIF-1α is recognised by the E3 ligase complex substrate recognition subunit, von Hippel-Lindau (VHL), leading to HIF-1α ubiquitination and subsequent proteasomal degradation^43^. Mitochondrial ROS, including RET-generated, can inhibit PHD and thereby stabilize HIF-1α, also under normoxic conditions^23,44–47^. To consolidate the role of HIF-1α signalling, we elicited transient stabilisation of HIF-1α in embryonic muscle DA1 at the control temperature of 25°C, via RNAi knockdown of HIF-1α degradation machinery components: the prolyl hydroxylase, PHD, or the VHL ubiquitin ligase complex substrate recognition subunit. Both manipulations phenocopy the characteristic critical period-induced NMJ overgrowth phenotype (Figure 4).

**Figure 4:**
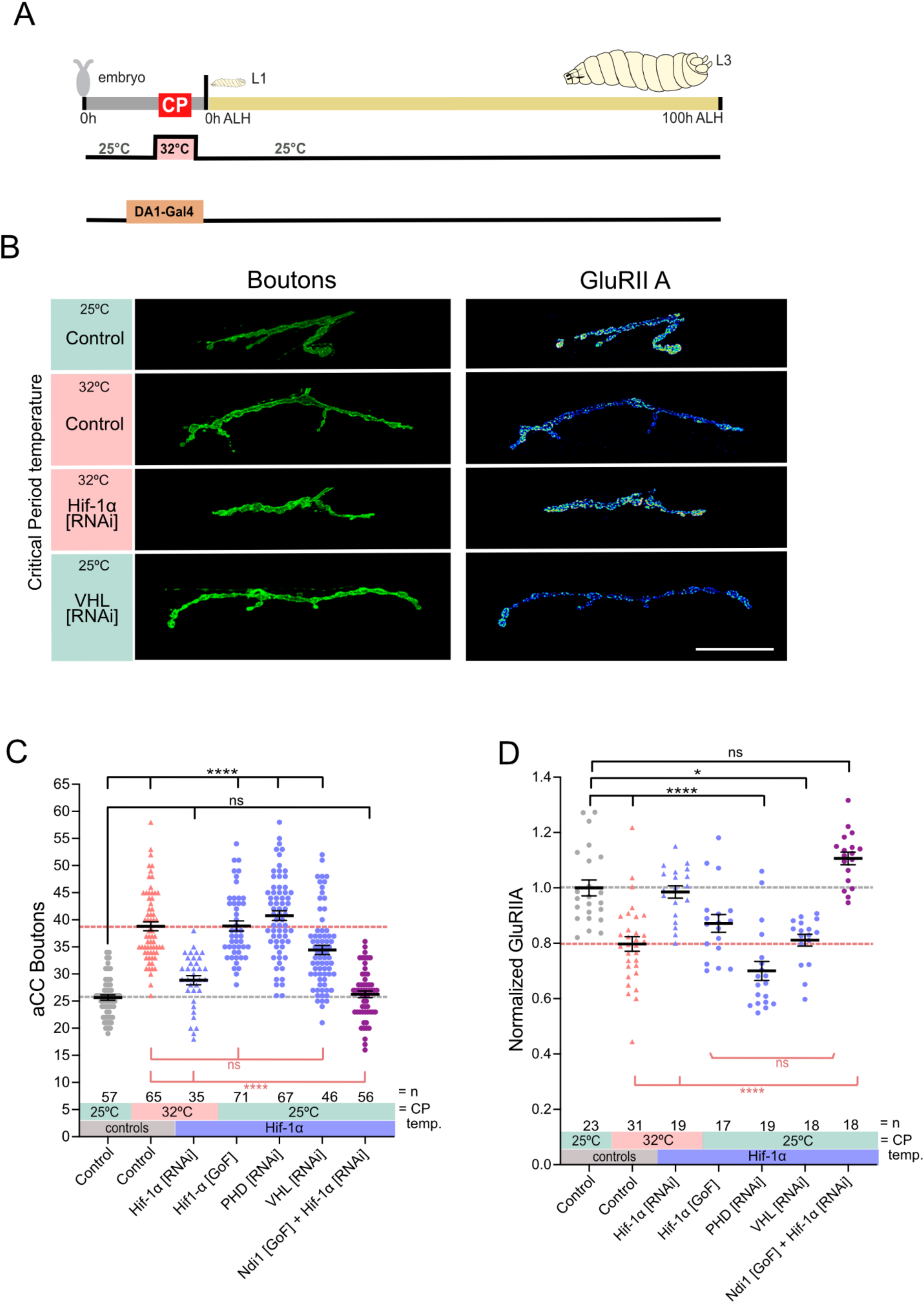
HIF-1α is the signal downstream of ROS-RET. **A)** Experimental paradigm **B)** Temperature experienced during the embryonic critical period (25°C control *vs* 32°C heat stress) and simultaneous genetic manipulation of muscle DA1 during embryonic stages only. “Control” indicates control genotype heterozygous for Oregon-R and DA1-Gal4. Larvae were reared at the control temperature of 25°C until the late wandering stage, 100 hrs ALH. GluRIIA and GluRIIB subunits are displayed with lookup table “fire” to illustrate signal intensities (warmer colours indicating greater signal intensities). Scale bar = 20 µm. **C)** Dot-plot quantification shows changes to aCC NMJ growth on its target muscle DA1, based on the standard measure of the number of boutons (swellings containing multiple presynaptic release sites/active zones). Data are shown with mean ± SEM, ANOVA, **** p<0.00001, ‘ns’ indicates statistical non-significance. Black asterisks indicate comparison with the control condition of 25°C throughout, genetically unmanipulated. Red asterisks indicate comparisons with control genotype exposed to 32°C heat stress during the embryonic critical period. **D)** Dot-plot quantification shows changes in levels of the GluRIIA receptor subunit at aCC NMJs quantified in **C)**. Data are shown with mean ± SEM, ANOVA, * p<0.01, **** p<0.00001, ‘ns’ indicates statistical non-significance. Black asterisks indicate comparison with the control condition of 25°C throughout, genetically unmanipulated. Red asterisks indicate comparisons with control genotype exposed to 32°C heat stress during the embryonic critical period. **E)** Dot-plot quantification as in **D)**, but for the low conductance GluRIIB receptor subunit, which remains unaffected by these manipulations.

To further validate a working model of HIF-1α signalling as downstream of mitochondrial RET-generated ROS, we used a genetic epistasis experiment: inducing RET via mis-expression of Ndi1, while simultaneously knocking down the putative downstream acceptor, HIF-1α. We found that RET induction was no longer able to induce the expected changes in the larval NMJ when HIF-1α was knocked down in the same cell (Figure 4). This supports a working model of information flow from mitochondrial RET-generated ROS to nuclear HIF-1α.

### Neuronal excitability and crawling speed are specified during the critical period by the mitochondrial ROS to HIF-1α signalling axis

Having thus far focused on the NMJ as a model synapse, we asked whether the structural changes that result from critical period plasticity have correlates in neuronal function and behaviour^48^. To this end, we performed whole cell patch clamp recordings from the aCC motoneuron at the late larval stage. These showed that embryonic heat stress of 32°C caused a lasting reduction in excitability (ie reduced capability to fire action potentials), while leaving cell size (assessed by membrane capacitance) and resting membrane potential unaffected (Figure 5A-D). Like NMJ critical period plasticity phenotypes, the effect on reducing neuronal excitability also requires the RET-ROS - HIF-1α signalling axis. It is blocked on neutralizing RET-ROS by expression of AOX. Conversely, induction of HIF-1α during the critical period is sufficient to induce the same reduction in aCC excitability at the control temperature of 25°C (Figure 5A-D).

**Figure 5.**
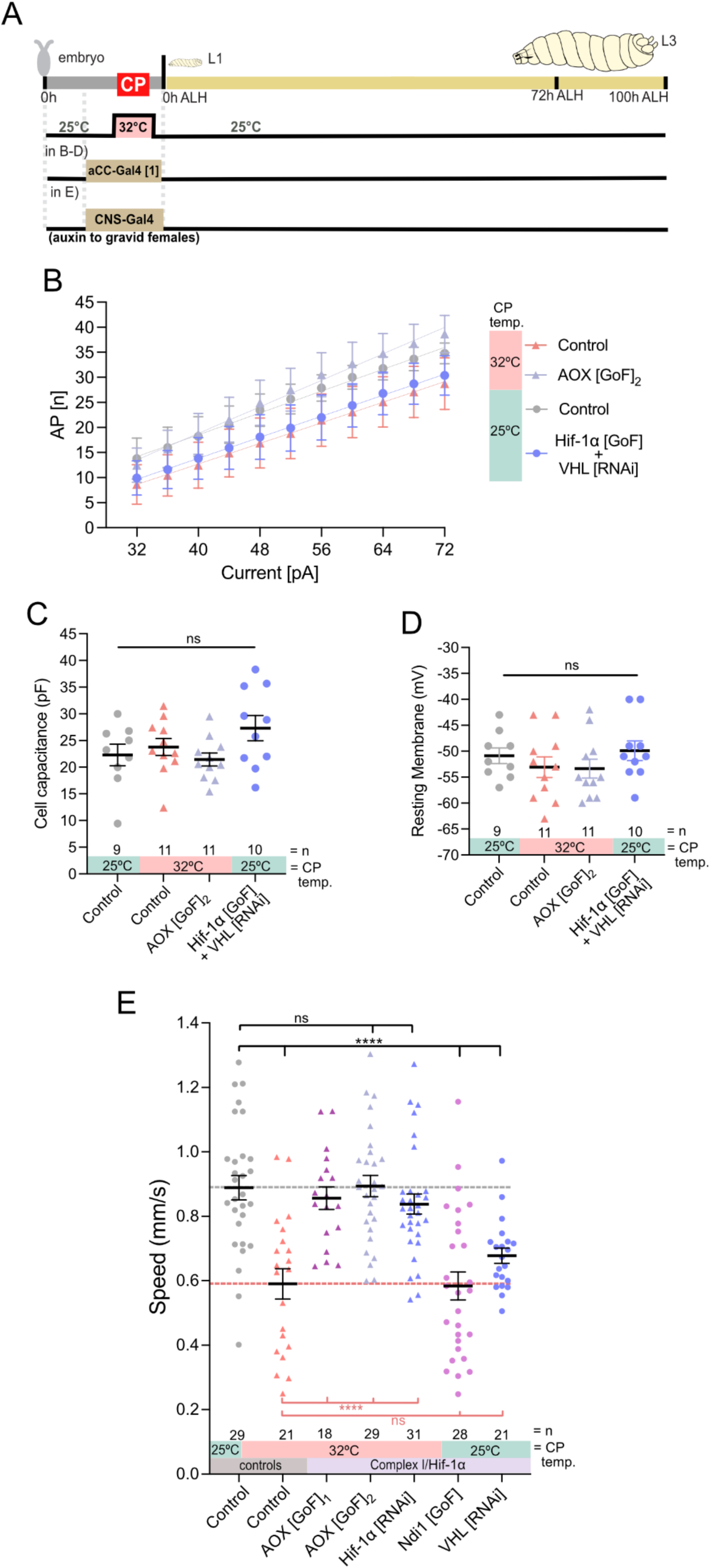
ROS by RET and HIF-1α signalling during the embryonic critical period are necessary and sufficient to reduce motoneuron excitability and crawling behaviour. **A)** Experimental paradigm **B)** aCC motoneuron excitability at the late third instar larval stage (i.e. action potential firing frequency triggered by current injected). Temperature experienced during the embryonic critical period (25°C control *vs* 32°C heat stress) and simultaneous transient genetic manipulation of aCC motoneuron during embryonic stages, via aCC-Gal4 [1] (RN2-O-Gal4). “Control” indicates control genotype heterozygous for Oregon-R and aCC-Gal4 [1]. Larvae were reared at the control temperature of 25°C until the late wandering stage, c. 100 hrs ALH. Control at 25°C *vs* 32°C is significant at p = 0.0005; Control at 25°C *vs* HIF-1α[GoF] is significant at p = 0.0015; Control at 25°C *vs* AOX[GoF]_2_ at 32°C is not significant at p = 0.28. **C)** Cell capacitance measurements as indicators of cell size show no significant differences between aCC motoneurons from specimens in B). **D)** Resting membrane potential, as an indicator for cell integrity, show no significant differences between aCC motoneurons from specimens in B). **E)** Crawling speed of third instar larvae (72 hrs ALH). Temperature experienced during the embryonic critical period (25°C control *vs* 32°C heat stress) and simultaneous genetic manipulation of CNS neurons during embryonic stages only. “Control” indicates control genotype heterozygous for Oregon-R and CNS-Gal4; Auxin-Gal80. Larvae were reared at the control temperature of 25°C. Each data point represents the crawling speed from an individual uninterrupted continuous forward crawl, n = specimen replicate number, up to three crawls assayed for each larva. Mean ± SEM, ANOVA, ns = not significant, ****p<0.00001.

At a network level, exposure to 32°C during the critical period results in reduced crawling speed at late larval stage (Figure 5E). We find that mitochondrial RET-ROS and HIF-1α signalling are required and sufficient for this critical period-regulated change in locomotor network output. Transient induction of these signals, in all embryonic neurons, at the control temperature, phenocopies the reduction in crawling speed caused by critical period heat stress (Figure 5E). RET-ROS and HIF-1α signalling only have this lasting effect when induced during the embryonic phase, including the critical period window, while manipulations outside this plastic period, i.e. post critical period closing, are without effect (supplementary figure 2A, B). Thus, the changes we observed, both centrally and at the NMJ, following critical period heat stress are sufficient to alter locomotor behaviour.

We considered two more likely scenarios to explain reduced locomotor speed. First, embryonic exposure to 32°C may lead to larvae that are simply unable to crawl as fast as control animals. Second, embryonic heat stress during the critical period may ‘set’ a different locomotor network homeostatic setpoint. To test this, we acutely increased the temperature during the behavioral crawling assay and found that larvae exposed to heat stress as embryos still responded with an increase in speed, as control animals do. Moreover, under such conditions they could match the crawling speed of controls reared at 25°C (Supplementary Figure 3A). However, when critical period-manipulated animals were maintained at 32°C for several hours, their crawling speed reverted to the original reduced level, suggesting that the system is still capable of homeostasis: responding and returning to a pre-determined “default velocity” (Supplementary Figure 3B). These observations suggest, that following critical period heat stress, the system is not simply damaged, but remains responsive to environmental temperature challenges, as in control animals. We speculate that following a critical period manipulation the locomotor network “default output” is positioned at a different set point or range.

We conclude that during the embryonic critical period, mitochondrial RET-generated ROS are instructive signals that are transmitted to the nucleus via the conserved transcriptional regulator, HIF-1α. Only when activated during the embryonic critical period, does this signalling pathway alter the subsequent development of the nervous system, leading to significant and permanent change in synaptic terminal growth, receptor composition, neuronal excitable properties and network output, which manifests as changes in larval crawling behaviour (Figure 6).

**Figure 6.**
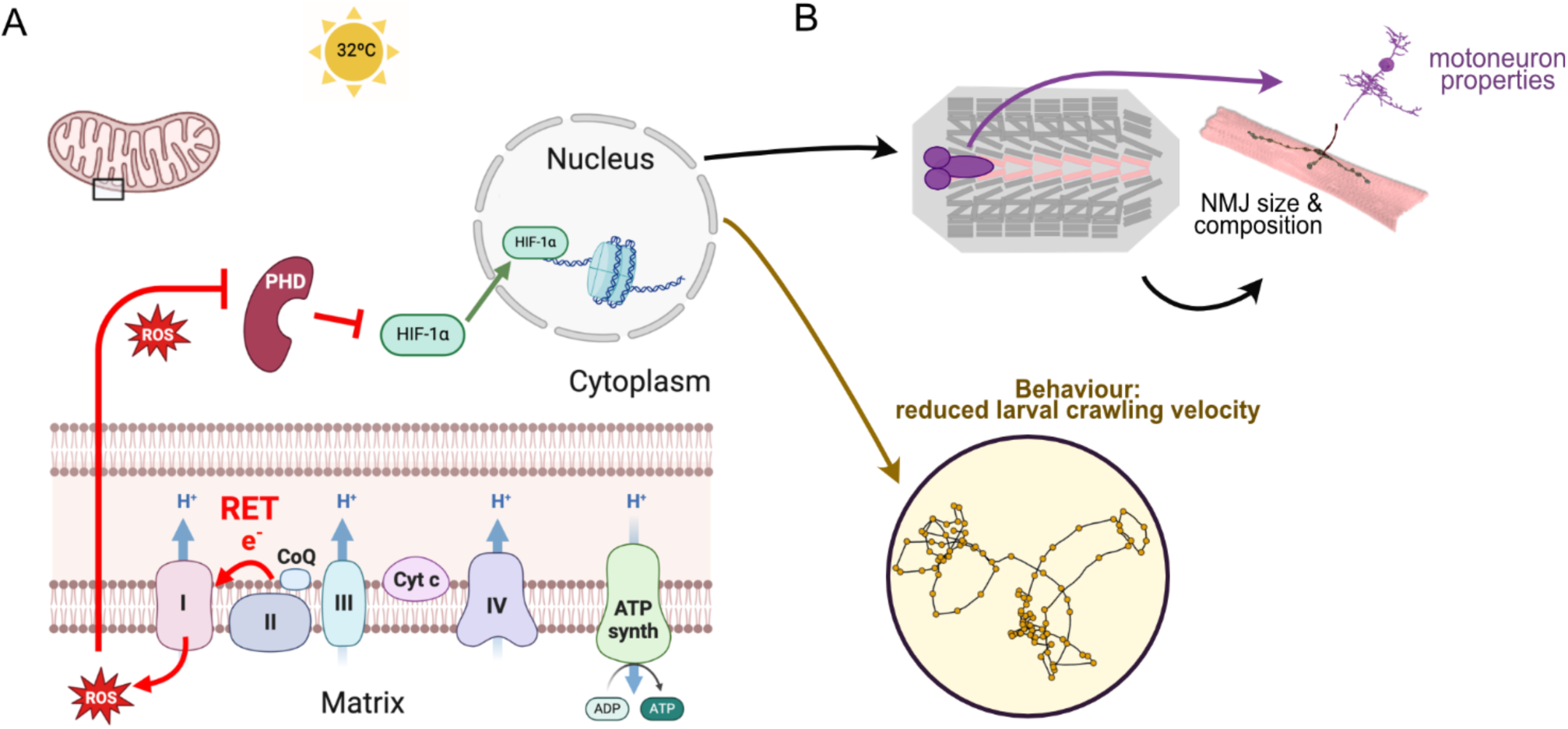
Working model. **A)** Heat stress during the critical period induces mitochondrial ROS, which lead to HIF-1α stabilisation and its accumulation in the nucleus, where HIF-1α generates long-lasting changes that impact on **B)** NMJ development: larger presynaptic terminals and less GluRIIA, as well as reductions in motoneuron excitability and, behaviourally, crawling speed.

## Discussion

Working with a simplified experimental system, we have identified a metabolic signal, mitochondrial RET-generated ROS, as a primary critical period plasticity signal, that is transduced to the nucleus via the conserved HIF-1α signalling pathway. Moreover, we demonstrate that critical period-plasticity responses are fundamentally cell intrinsic, shown by virtue of targeting genetic manipulations, transiently, to single cells *in vivo*. This provides a first mechanistic explanation for how a perturbation experienced during a critical period effects a change in subsequent development, leading to altered neuronal properties and animal behaviour.

### Environmental cues are integrated during critical periods and direct nervous system development

It has long been known that early life experiences are formative, due to their potential to impart lasting influence on nervous system function^49,50^. Though critical periods have been most intensively studied in the context of activity-regulated sensory processing and cortical networks, it is possible that critical periods represent a fundamental process that is common to the development of nervous systems. For example, similar developmental windows of heightened vulnerability to disturbances have also been identified in mammalian locomotor systems^51^, as well as in locomotor, visual and olfactory systems of zebrafish^52–56^ and several insect species (ants, bees and *Drosophila*^57,57–61)^. In fast developing organisms, opening of a critical period correlates with the phase when network activity transitions from spontaneous and un-patterned to patterned activity, and in the case of the locomotor network of the *Drosophila* embryo lasts only two hours^11,13,14,62–64^. In mammalian systems, critical periods are much more drawn out, lasting weeks in mouse and up to several years in human^7^. Notwithstanding such differences, the neuronal activity-regulated processes of critical periods appear remarkably similar across species; from GABAergic signalling during critical periods of visual system and cortex development^56^, to the importance of networks attaining an appropriate excitation:inhibition balance, also recently shown for the developing *Drosophila* locomotor network^12,13^.

Here, we took advantage of the well-defined critical period of the *Drosophila* locomotor network^11,12^. As an ecologically relevant stimulus, we focused on temperature, specifically a 32°C heat stress, which we have previously shown phenocopies pharmacological or genetic activity manipulations, leading to similar lasting changes in subsequent neuronal development^48^. Using this experimental paradigm allowed us to identify a basic instructive signal that underlies critical period-plasticity, namely mitochondrial ROS generated at Complex-I via reverse flow of electrons within the electron transport chain. In mammals, mitochondrial RET-generated ROS has been associated with ageing and expression of protective responses^19,65^. Whether this same signal is also responsible for the responses to developmental heat stress reported in ants^66^, bees^59,67,68^ or other parts and stages of the developing *Drosophila* nervous system^15,57^, remains to be tested. Though we employed a stress signal in this study, to elicit a robust critical period plasticity response, temperature operates as a general environmental cue during nervous system development. Complementing this study, a cooler, non-stress temperature experienced during a pupal critical period of *Drosophila* nervous system development, has been reported to also lead to clear cellular effects. In that case, causing reduced filopodial dynamics and a consequent increase in synapse formation with concomitant changes in network connectivity and behavioural output^57,61^.

### Critical period effects at the level of single cells

One of the strengths of the experimental system we use is that it allows studying critical periods with single cell resolution *in vivo*. Here, we demonstrate that critical period plasticity is primarily cell autonomous. This simplifies establishing clear cause-effect relationships, which is more difficult with commonly used network-wide manipulations. Through cell-selective genetic manipulations of a single motoneuron or its target muscle, we phenotypically rescue individual cells from systemic heat stress manipulations. Thus, we demonstrate the cell-autonomous necessity of RET-generated mitochondrial ROS and HIF-1α signalling for heat stress-induced critical period plasticity. Conversely, we also show cell-selective sufficiency of RET-ROS and HIF-1α signalling to phenocopy changes in NMJ development, though only when induced during the transient embryonic critical period, but not after.

From a developmental perspective, this implies that different cell types might be integrated into circuits in a sequential manner, for example based on birth order or maturation speed, thus facilitating more reproducible and robust network assembly. Across a developing nervous system, different regions are known to undergo their respective critical periods in sequence^7^, which may be mirrored at the cellular level within a given network. A recent study centred on developing mouse neocortex demonstrated the sequential maturation of two principal classes of inhibitory interneurons, and how this process directs network maturation^69^.

### Is the critical period a developmental window during which network setpoints are established?

Neuronal networks dynamically maintain stability and appropriate function around a given set point (or range), through homeostatic plasticity mechanisms^70,71^. How and when a set point is specified remains unknown. Previous evidence suggests that perturbations of activity during a critical period can alter the homeostatic set point, and thus lead to long-lasting changes in network function, including instability and epilepsy-like activity^11,72,73^. Thus, the focus has been on neuronal activity and its coordination as important processes in setting the set point^11,12,62–64,74,75^. A challenge has been to relate the consequences of perturbing network activity, during a critical period, to specific changes in functional or behavioural outcomes. By working with a locomotor network, we found that heat stress at the cellular level, as a critical period perturbation, leads to reduced excitability of motoneurons. At the network level, this correlates with reduced crawling speed, which is seen as directly reflecting a change in output of the locomotor network. Importantly, changes in crawling speed only occurred following manipulations that spanned the embryonic critical period, and animals maintained the ability to increase their crawling speed. Moreover, network homeostasis appears to remain intact, as critical period-manipulated “slow” animals respond normally to acute upshifts in environmental temperature, but transform back to their slower “default” speed following chronic changes^30^. These observations are compatible with the possibility that the set point for locomotor network output is established during the critical period. Because this network is so well characterised and experimentally tractable, it now offers the opportunity to study the cellular mechanisms through which this is achieved.

### Conserved mitochondrial-nuclear signalling directs critical period plasticity

Cells integrate internal (e.g. network activity) and external environmental cues (e.g. temperature) during a critical period^12^. The identification of an instructive signal that is intimately linked to mitochondrial metabolism significantly adds to, and complements, a large body of work that has focused on activity-regulated processes^1,2^. How a neuronal activity-regulated signal and mitochondrial metabolism interlink, especially during critical periods of development, will be important to understand. We had previously shown that mitochondrial and NADPH oxidase-generated ROS are required for activity-regulated neuronal plasticity outside the critical period^30,31,76^. Here, we identified a specific source of mitochondrial ROS, namely those generated at Complex-I via RET, which occurs when the coenzyme Q pool is reduced while the proton motif force is elevated^77^. A mitochondrial enzyme that is linked to the redox state of the coenzyme Q pool, dihydroorotate dehydrogenase (DHODH), has been identified as an important regulator of neuronal action potential firing rate, i.e. a determinant of the neuronal homeostatic setpoint^78,79^. Together with our findings here, we speculate that mitochondrial metabolism might be central to critical period plasticity, potentially encoding neuronal homeostatic setpoints.

How might a transient perturbation during a critical period lead to lasting change? HIF-1α, a transcriptional regulator involved in the cellular response to stress, is typically degraded under normal oxygen concentrations. The model we propose is that mitochondrial ROS, including those generated by RET, transiently inhibit the degradation of HIF-1α, thus allowing HIF-1α to accumulate and translocate into the nucleus^23,23,44–47^. HIF-1α signalling is a known regulator of changes in cellular metabolism, for example promoting aerobic glycolysis in a range of cancers^80–82^, but also during normal development, when rapid tissue growth is required^26,83,84^. HIF-1α achieves this through a combination of direct regulation of gene expression, notably of metabolic enzymes, and epigenetic modifications, partly executed through histone acetyl transferases that act as co-factors^85–88^. In some forms of cancer, this is thought to create positive feedback loops, from elevated mitochondrial metabolism, to increased ROS production maintaining disinhibition of HIF-1α, which reinforces metabolic drive. It is therefore conceivable that transient HIF-1α signalling could initiate such metabolic-epigenetic feedback loops that maintain a change in metabolic state, and thus also of neuronal growth and functional properties.

## Materials and Methods

### Drosophila rearing and stocks

Drosophila stocks were kept on standard corn meal medium, at 25°C. For all experiments where crosses were necessary, stocks were selected and combined as a 1:2 ratio of Gal4-line males to UAS-line virgin females. See Table 1 for the strains used.

**Table 1.** Drosophila strains.

| Drosophila strain name | Additional information | Reference |
| --- | --- | --- |
| Oregon-R-P2 | Wild Type | Bloomington Drosophil Stock Center; RRID:BDSC_2376 |
| eve-Gal4<sup>eme</sup> (aka SS2-C-Gal4) | Gal4 expression targeted to muscle DA1 during embryonic stages only | [90] |
| aka RN2-O-gal4 (aCC-Gal4 [1]) | Gal4 expression targeted to motoneurons aCC and RP2 during embryonic stages only | [91] |
| GMR94G06-Gal4 (aCC-Gal4 [2]) | Gal4 expression targeted to motoneuron aCC | Bloomington Drosophil Stock Center; RRID:BDSC_40701 |
| GMR57C10-GAL4 (n-synaptobrevin-Gal4) | Gal4 expression targeted to neurons | Bloomington Drosophil Stock Center; RRID:BDSC_39171 |
| AtTIR1-T2A-AID-GAL80-AID (Auxin-gal80) | Ubiquitously express GAL80 sensitive to auxin | [41]; gift from Tony Southall |
| UAS-mito-catalase | Mitochondrial targeted catalase under UAS control | gift from Alex Whitworth |
| UAS-Prx3 | Peroxiredoxin 3 under UAS control | gift from Alberto Sanz |
| UAS-ND75 [RNAi] | Expresses dsRNA for RNAi of ND75 (FBgn0017566) under UAS control | Vienna Drosophila Resource Center #100733/KK; gift from Alberto Sanz |
| UAS-AOX [GoF]1 | Expresses AOX from C. intestinalis under UAS control (allele F6) | [19]; gift from Alberto Sanz |
| UAS-AOX [GoF]2 | Expresses AOX from C. intestinalis under UAS control (allele F24) | [19]; gift from Alberto Sanz |
| UAS-Ndi1 [GoF] | Expresses Ndi1 from S.cerevisiae under UAS control (allele A46 and B20) | [20]; gift from Alberto Sanz |
| UAS-Sima [RNAi] | Expresses dsRNA for RNAi of sima (FBgn0266411) under UAS control | Bloomington Drosophil Stock Center; RRID:BDSC_26207 |
| UAS-Sima [GoF] | Expresses sima (FBgn0266411) under UAS control | Gift from Joe Bateman |
| UAS-VHL [RNAi] | Expresses dsRNA for RNAi of VHL (FBgn0041174) under UAS control | Bloomington Drosophil Stock Center; RRID:BDSC_50727 |
| UAS-PHD [RNAi] | Expresses dsRNA for RNAi of PHD (FBgn0264785) under UAS control | Bloomington Drosophil Stock Center; RRID:BDSC_34717 |

### Critical Period manipulations

#### Temperature manipulations

Embryos were harvested from 0 to 6 hours after egg lay at 25°C. Embryos were then shifted to their respective temperatures in incubators (25°C or 32°C) for the remainder of embryogenesis. After all animals hatched, all were moved back to 25°C until assayed in late larval stages.

#### Auxin feeding

Auxin (GK2088 1-Naphthaleneacetic acid potassium salt; Glentham) solution was mixed with melted molasses fly medium to achieve 5mM auxin concentration and poured into petri dish plates. For embryonic exposure to auxin, adult flies in mixed-sex laying pots were fed on this food for 48 hours prior to egg collection. For larval exposure, freshly hatched larvae were fed on this food for 24 hours.

### Dissections and immunocytochemistry

Flies were allowed to lay eggs on apple-juice agar based medium overnight at 25 °C. Larvae were then reared at 25°C on yeast paste, while avoiding over-crowding. Precise staging of the late wandering third instar stage was achieved by: a) checking that a proportion of animals from the same time-restricted egg lay had initiated pupariation; b) larvae had reached a certain size and c) showed gut-clearance of food (yeast paste supplemented with Bromophenol Blue Sodium Salt (Sigma-Aldrich)). Larvae (of either sex) were dissected in Sorensen’s saline, fixed for 10 min at room temperature in Bouins fixative (Sigma-Aldrich), as previously detailed^31^. Wash solution was Sorensen’s saline containing 0.3% Triton X-100 (Sigma-Aldrich) and 0.25% BSA (Sigma-Aldrich). Primary antibodies, incubated overnight at 10°C, were: Goat-anti-HRP Alexa Fluor 488 (1:1000; Jackson ImmunoResearch Cat. No. 123-545-021), Rabbit-anti-GluRIIB (1:1000; gift from Mihaela Serpe^89^), Mouse anti-GluRIIA (1:100; Developmental Studies Hybridoma Bank, Cat. No. 8B4D2 (MH2B)); Mouse anti-BRP (1:100; Developmental Studies Hybridoma Bank, Cat. No. nc82); secondary antibodies, 2 hr at room temperature: Donkey anti-Mouse StarRed (1:1000; Abberior, cat.no. STRED-1001); and goat anti-Rabbit Atto594 (1:1000; Sigma-Aldrich Cat No 77671-1ML-F). Specimens were cleared in 80% glycerol, overnight at 4°C, then mounted in Mowiol.

### Image acquisition and analysis

Specimens were imaged using an Olympus FV3000 point-scanning confocal, and a 60x/1.3 N.A silicone oil immersion objective lens and FV31S-SW software. Confocal images were processed using ImageJ (to quantify GluRIIA and -B intensity) and Affinity Photo (Serif Ltd) to prepare figures. Bouton number of the aCC NMJ on muscle DA1 from segments A3-A5 was determined by counting every distinct spherical varicosity.

To quantify GluRIIA and -B fluorescence intensity a threshold image for each channel was used to generate an outline selection, which included only the postsynaptic glutamate receptor clusters. This outline was then applied to the original image to define the regions of interest for analysis. Fluorescence intensities for GluRIIA and GluRIIB channels was measured within these outlines. Data were exported into Excel and the intensity values normalised to the negative control condition (25°C through all embryogenesis).

### Electrophysiology

Third instar (L3) larvae (of either sex) were dissected in a dish containing standard saline (135mM NaCl (Fisher Scientific), 5mM KCl (Fisher Scientific), 4mM MgCl2·6H2O (Sigma-Aldrich), 2mM CaCl2·2H2O (Fisher Scientific), 5mM TES (Sigma-Aldrich), 36mM sucrose (Fisher Scientific), pH 7.15) to remove the ventral nerve cord and brain lobes (CNS). The isolated CNS was then transferred to a droplet of external saline containing 200 µM mecamylamine (Sigma-Aldrich) to block postsynaptic nACh receptors in order to synaptically isolate motor neurons. CNSs were laid flat (dorsal side up) and glued (GLUture Topical Tissue Adhesive; World Precision Instruments USA) to a Sylgard-coated cover slip (1 to 2mm depth of cured SYLGARD Elastomer (Dow-Corning USA) on a 22 x 22mm square coverslip). Preparations were placed on a glass slide under a microscope (Olympus BX51-WI), viewed using a 60x water-dipping lens. To access nerve cell somata, 1% protease (Streptomyces griseus, Type XIV, Sigma-Aldrich, in external saline) contained within a wide-bore glass pipette (GC100TF-10; Harvard Apparatus UK, approx. 10 µm opening) was applied to abdominal segments, roughly between A5-A2 (Baines and Bate, 1998) to remove overlaying glia. Motoneurons were identified by anatomical position and relative cell size, with aCC being positioned close to the midline and containing both an ipsilateral and contralateral projection. Whole cell patch clamp recordings were made using borosilicate glass pipettes (GC100F-10, Harvard Apparatus) that were fire polished to resistances of 10 -15MΩ when filled with intracellular saline (140 mM potassium-D-gluconate (Sigma-Aldrich), 2 mM MgCl2·6H2O (Sigma-Aldrich), 2 mM EGTA (Sigma-Aldrich), 5 mM KCl (Fisher Scientific), and 20 mM HEPES (Sigma-Aldrich), (pH 7.4). Input resistance was measured in the ‘whole cell’ configuration, and only cells that had an input resistance ≥ 0.5 GΩ were used for experiments. Cell capacitance and break-in resting membrane potential were measured for each cell recorded; only cells with a membrane potential upon break in of < - 40 mV were analysed. Data for current step recordings were captured using a Multiclamp 700B amplifier (Molecular Devices) controlled by pCLAMP (version 10.7.0.3), via an analogue-to-digital converter (Digidata 1440A, Molecular Devices). Recordings were sampled at 20 kHz and filtered online at 10 kHz. Once patched, neurons were brought to a membrane potential of -60 mV using current injection. Each recording consisted of 20 x 4 pA (500 ms) current steps, including an initial negative step, giving a range of -4 to +72 pA. Number of action potentials were counted at each step and plotted against injected current. Cell capacitance and membrane potential were measured between conditions to ensure that any observed differences in excitability were not due to differences in either cell size or resting state.

### Larval crawling analysis

At the mid-third instar stage, 72 hours after larval hatching (ALH), larvae were rinsed in water and placed inside a 24cm x 24cm crawling arena with a base of 5mm thick 0.8% agarose gel (Bacto Agar), situated inside an incubator. Temperature was maintained at 25±0.5°C, reported via a temperature probe in the agar medium. Humidity was kept constant. Larval crawling was recorded using a frustrated total internal reflection-based imaging method (FIM) in conjunction with the tracking software FIMTrack (Risse et al., 2017, 2013) using a Basler acA2040-180km CMOS camera with a 16mm KOWA-IJM3sHC.SW-VIS-NIR Lens controlled by Pylon (by Basler) and Streampix (v.6) software (by NorPix). Larvae were recorded for 20 minutes at five frames per second.

Recordings were split into four 5-minute sections with the first section not tracked and analysed as larvae are acclimatising to the crawling arena. The remaining three sections were used to analyse crawling speed, with each larva sampled at most once per section. Crawling speed was calculated using FIMTrack software choosing crawling tracks that showed periods of uninterrupted forward crawling devoid of pauses, turning or collision events. The distance travelled was determined via the FIMTrack output.

### Statistical analysis

Statistical analyses were carried out using GraphPad Prism Software (Version 10.1.2). Datasets were tested for normal distribution with the Shapiro-Wilk Test. Normally distributed data were then tested with students t-test (for pairwise comparison). Normally distributed analysis for more than two groups was done with a one-way ANOVA and post hoc tested with a Tukey multiple comparison test. Non-normally distributed data sets of two groups were tested with Mann-Whitney U Test (pairwise comparison) and datasets with more than two groups were tested with a Kruskal Wallis ANOVA and post hoc tested with a Dunns multiple post hoc comparison test. For whole cell electrophysiology in aCC neurons, linear regression analysis was used to compare the intercepts between conditions, using a Bonferoni correction for multiple comparisons, resulting in an adjusted significance threshold of p < 0.0083. This analysis was restricted to the linear portions of the input-output curves. For all datasets mean and standard error of mean (SEM) are shown. Significance levels were * p<0.05; ** p<0.01; *** p<0.001; **** p<0.0001.

## Author Contributions

Conceptualization: D.S.-C. and M.L; Investigation: D.S.-C., B.C., M.M., M.C.W.O. and T.P.; Writing – Original Draft: D.S.-C. and M.L; Writing – Review & Editing: D.S.-C., B.C, M.C.W.O., T.P., R.A.B. and M.L.; Supervision: R.A.B and M.L.; Funding Acquisition: D.S-C., R.A.B and M.L.

## Funding

D.S.-C. was supported by the European Molecular Biology Organization (EMBO) with a long-term EMBO fellowship (ALTF 62-2021). This work was supported by a Joint Wellcome Trust Investigator Award to R.A.B. and M.L. (217099/Z/19/Z).

### Acknowledgments

The authors are grateful to Mihaela Serpe for the Rabbit polyclonal anti-GluRIIB antiserum. The authors are grateful to Tony Southall, Alex Whitworth, Joe Bateman and Alberto Sanz for generously providing fly stocks. Stocks obtained from the Bloomington Drosophila Stock Center (NIH P40OD018537) were used in this study. The nc82 monoclonal antibody was obtained from the Developmental Studies Hybridoma Bank, created by the NICHD of the NIH and maintained at The University of Iowa, Department of Biology, Iowa City, IA 52242. The work benefited from the Imaging Facility, Department of Zoology, supported by Matt Wayland, and funds from a Wellcome Trust Equipment Grant (WT079204) with contributions by the Sir Isaac Newton Trust in Cambridge, including Research Grant [18.07ii(c)]. Work on this project benefited from the Manchester Fly Facility, established through funds from the University of Manchester and the Wellcome Trust (Grant 087742/Z/08/Z).

## Supplementary Material

**Supplementary figure 1.**
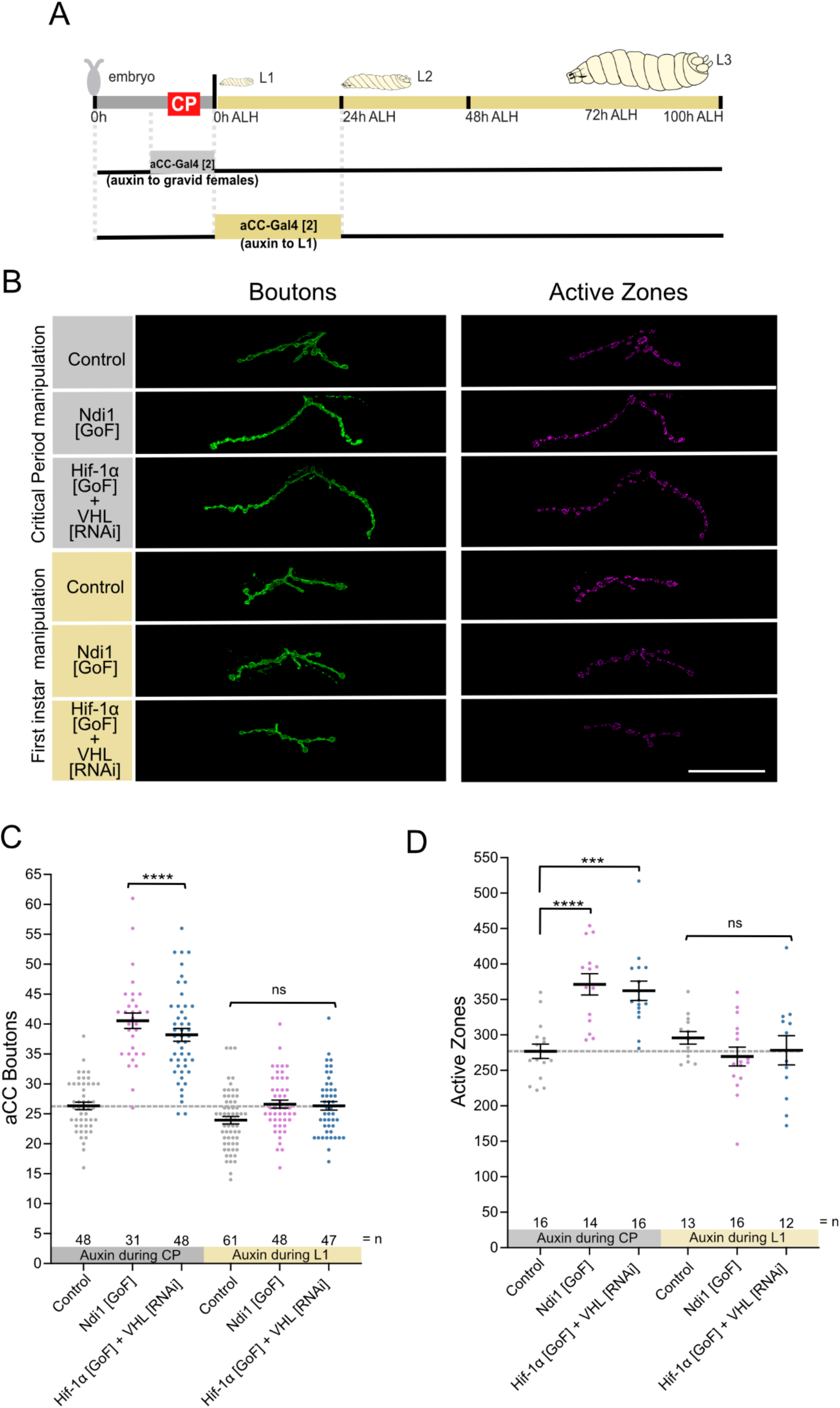
RET-ROS and HIF-1α signalling has lasting impact on NMJ only during the Critical Period. **A)** Experimental paradigm **B)** Genetic manipulation of motoneuron aCC during the critical period *vs* after critical period closure. “Control” indicates control genotype heterozygous for Oregon-R and aCC-Gal4[2]; Auxin-Gal80. Larvae were reared at the control temperature of 25°C until the late wandering stage, 100 hrs ALH. Scale bar = 20 µm. **C)** Dot-plot quantification shows changes to aCC NMJ growth on its target muscle DA1, based on the standard measure of the number of boutons (swellings containing multiple presynaptic release sites/active zones). Data are shown with mean ± SEM, ANOVA, **** p<0.00001, ‘ns’ indicates statistical non-significance. **D)** Dot-plot quantification shows changes in number of the active zones at aCC NMJs quantified in C). Data are shown with mean ± SEM, ANOVA, *** p<0.0001, **** p<0.00001, ‘ns’ indicates statistical non-significance.

**Supplementary figure 2.**
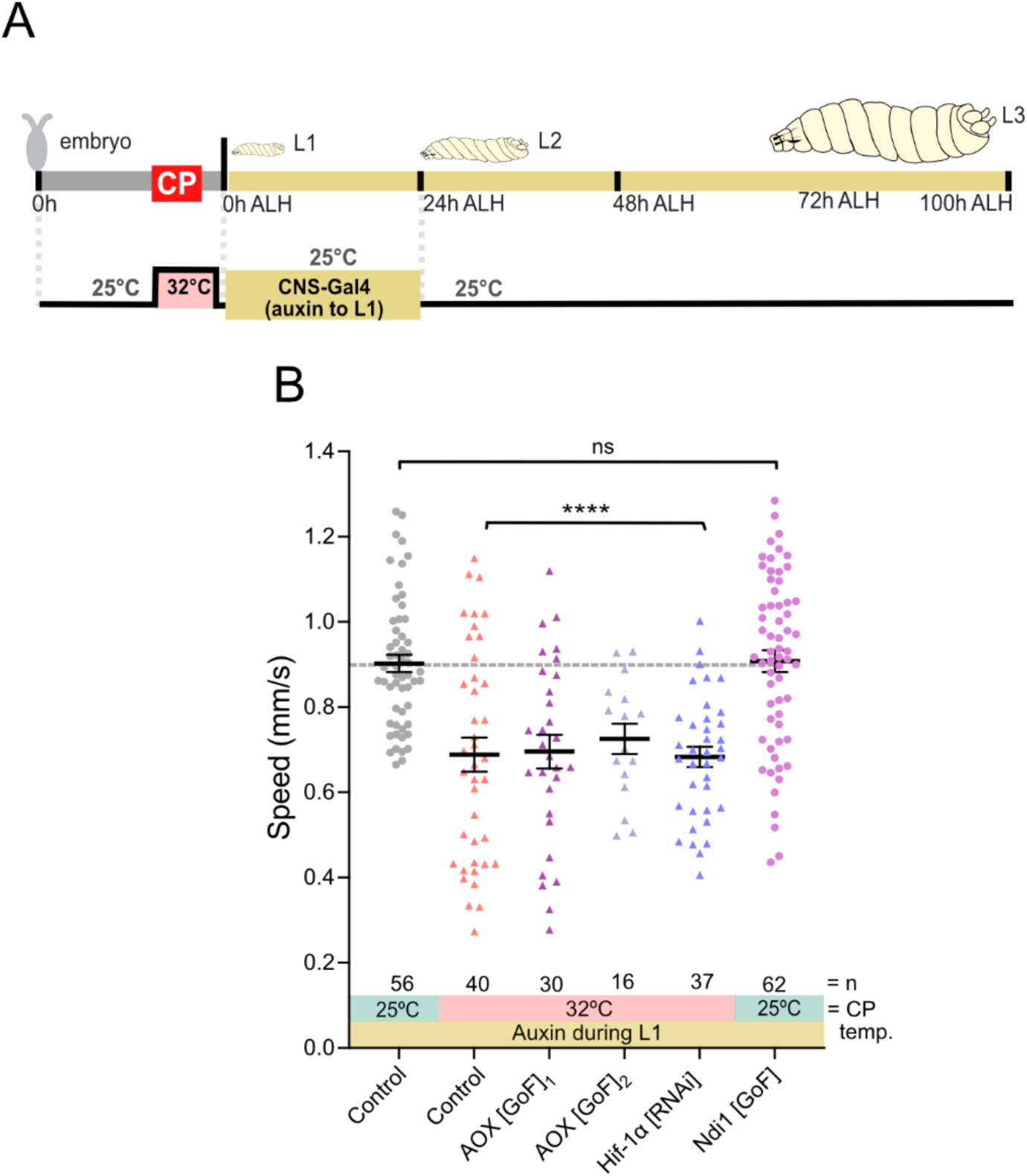
RET-ROS and HIF-1α signalling has not lasting impact on behaviour after the Critical Period. **A)** Experimental paradigm **B)** Crawling speed of third instar larvae (72 hrs after larval hatching (ALH)). Temperature experienced during the embryonic critical period (25°C control *vs* 32°C heat stress) and simultaneous genetic manipulation of all neurons during first instar stage. “Control” indicates control genotype heterozygous for Oregon-R and CNS-Gal4; Auxin-Gal80. Larvae were reared at the control temperature of 25°C until 72 hrs ALH. Each data point represents crawling speed from an individual uninterrupted continuous forward crawl, n = specimen replicate number, up to three crawls assayed for each larva. Mean ± SEM, ANOVA, ns = not significant, ****p<0.00001.

**Supplementary figure 3.**
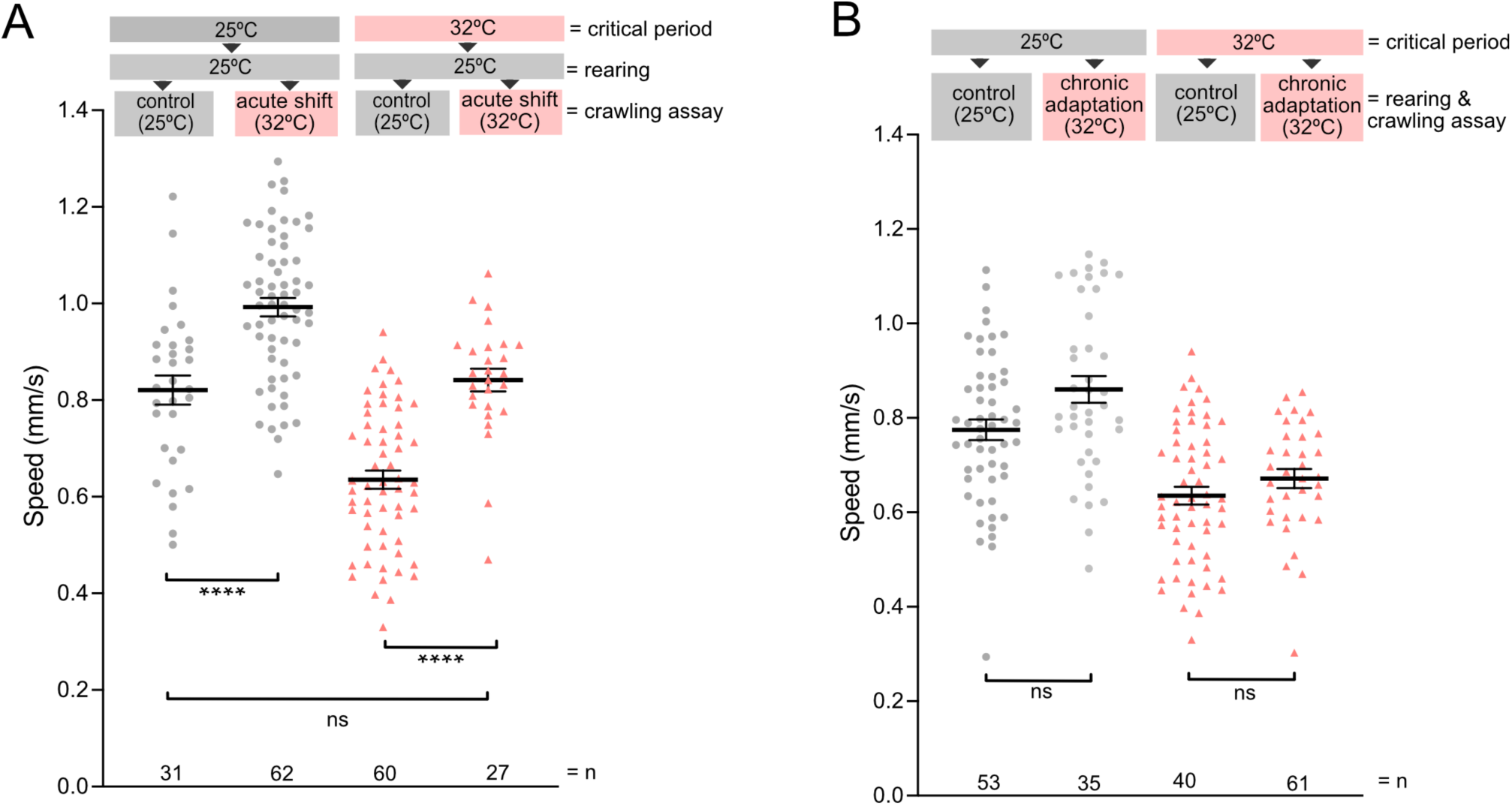
Homeostatic responses to environmental change are maintained after 32°C critical period. **A)** Acute shift of temperature. Crawling speed of third instar larvae (72 hrs after larval hatching (ALH)). Temperature experienced during the embryonic critical period (25°C control *vs* 32°C heat stress); temperature experienced during the larva rearing (25°C control *vs* 32°C heat stress) and control (25°C) versus acute shift of temperature during the crawling assay (32°C). Genotype is Oregon-R. Each data point represents crawling speed from an individual uninterrupted continuous forward crawl, n = specimen replicate number, up to three crawls assayed for each larva. Mean ± SEM, ANOVA, ns = not significant, ****p<0.00001. **B)** Chronic adaptation to the heat stress. Crawling speed of third instar larvae (72 hrs after larval hatching (ALH)). Temperature experienced during the embryonic critical period (25°C control vs 32°C heat stress); temperature experienced during the larva rearing and during the crawling assay (25°C control vs 32°C chronic adaptation). Oregon-R. Each data point represents crawling speed from an individual uninterrupted continuous forward crawl, n = specimen replicate number, up to three crawls assayed for each larva. Mean ± SEM, ANOVA, ns = not significant

## References

1. Hensch, T.K. (2004). CRITICAL PERIOD REGULATION. Annu. Rev. Neurosci. 27, 549–579. 10.1146/annurev.neuro.27.070203.144327.

2. Hensch, T.K. (2005). Critical period plasticity in local cortical circuits. Nat. Rev. Neurosci. 6, 877–888. 10.1038/nrn1787.

3. Hensch, T.K., and Quinlan, E.M. (2018). Critical periods in amblyopia. Vis. Neurosci. 35, E014. 10.1017/S0952523817000219.

4. Do, K.Q., Cuenod, M., and Hensch, T.K. (2015). Targeting Oxidative Stress and Aberrant Critical Period Plasticity in the Developmental Trajectory to Schizophrenia. Schizophr. Bull. 41, 835–846. 10.1093/schbul/sbv065.

5. Steullet, P., Cabungcal, J.-H., Coyle, J., Didriksen, M., Gill, K., Grace, A.A., Hensch, T.K., LaMantia, A.-S., Lindemann, L., Maynard, T.M., et al. (2017). Oxidative stress-driven parvalbumin interneuron impairment as a common mechanism in models of schizophrenia. Mol. Psychiatry 22, 936–943. 10.1038/mp.2017.47.

6. Sakai, A., and Sugiyama, S. (2018). Experience-dependent transcriptional regulation in juvenile brain development. Dev. Growth Differ. 60, 473–482. 10.1111/dgd.12571.

7. Reh, R.K., Dias, B.G., Nelson, C.A., Kaufer, D., Werker, J.F., Kolb, B., Levine, J.D., and Hensch, T.K. (2020). Critical period regulation across multiple timescales. Proc. Natl. Acad. Sci. 117, 23242–23251. 10.1073/pnas.1820836117.

8. Collins, C.A., Wairkar, Y.P., Johnson, S.L., and DiAntonio, A. (2006). Highwire Restrains Synaptic Growth by Attenuating a MAP Kinase Signal. Neuron 51, 57–69. 10.1016/j.neuron.2006.05.026.

9. Shen, W., and Ganetzky, B. (2009). Autophagy promotes synapse development in *Drosophila*. J. Cell Biol. 187, 71–79. 10.1083/jcb.200907109.

10. Frank, C.A., Wang, X., Collins, C.A., Rodal, A.A., Yuan, Q., Verstreken, P., and Dickman, D.K. (2013). New Approaches for Studying Synaptic Development, Function, and Plasticity Using *Drosophila* as a Model System. J. Neurosci. 33, 17560–17568. 10.1523/JNEUROSCI.3261-13.2013.

11. Giachello, C.N.G., and Baines, R.A. (2015). Inappropriate Neural Activity during a Sensitive Period in Embryogenesis Results in Persistent Seizure-like Behavior. Curr. Biol. 25, 2964–2968. 10.1016/j.cub.2015.09.040.

12. Hunter, I., Coulson, B., Pettini, T., Davies, J.J., Parkin, J., Landgraf, M., and Baines, R.A. (2024). Balance of activity during a critical period tunes a developing network. eLife 12, RP91599. 10.7554/eLife.91599.

13. Carreira-Rosario, A., York, R.A., Choi, M., Doe, C.Q., and Clandinin, T.R. (2021). Mechanosensory input during circuit formation shapes Drosophila motor behavior through patterned spontaneous network activity. Curr. Biol. 31, 5341–5349.e4. 10.1016/j.cub.2021.08.022.

14. Zeng, X., Komanome, Y., Kawasaki, T., Inada, K., Jonaitis, J., Pulver, S.R., Kazama, H., and Nose, A. (2021). An electrically coupled pioneer circuit enables motor development via proprioceptive feedback in Drosophila embryos. Curr. Biol. 31, 5327–5340.e5. 10.1016/j.cub.2021.10.005.

15. Williamson, M., Mitchell, A., and Condron, B. (2021). Birth temperature followed by a visual critical period determines cooperative group membership. J. Comp. Physiol. A 207, 739–746. 10.1007/s00359-021-01512-3.

16. Somero, G.N., Lockwood, B.L., and Tomanek, L. (2017). Biochemical adaptation: response to environmental challenges from life’s origins to the Anthropocene (Sinauer associates, Inc. publishers).

17. Scialò, F., Sriram, A., Fernández-Ayala, D., Gubina, N., Lõhmus, M., Nelson, G., Logan, A., Cooper, H.M., Navas, P., Enríquez, J.A., et al. (2016). Mitochondrial ROS Produced via Reverse Electron Transport Extend Animal Lifespan. Cell Metab. 23, 725–734. 10.1016/j.cmet.2016.03.009.

18. Scialò, F., Fernández-Ayala, D.J., and Sanz, A. (2017). Role of Mitochondrial Reverse Electron Transport in ROS Signaling: Potential Roles in Health and Disease. Front. Physiol. 8, 428. 10.3389/fphys.2017.00428.

19. Scialò, F., Sriram, A., Stefanatos, R., Spriggs, R.V., Loh, S.H.Y., Martins, L.M., and Sanz, A. (2020). Mitochondrial complex I derived ROS regulate stress adaptation in Drosophila melanogaster. Redox Biol. 32, 101450. 10.1016/j.redox.2020.101450.

20. Sanz, A., Soikkeli, M., Portero-Otín, M., Wilson, A., Kemppainen, E., McIlroy, G., Ellilä, S., Kemppainen, K.K., Tuomela, T., Lakanmaa, M., et al. (2010). Expression of the yeast NADH dehydrogenase Ndi1 in *Drosophila* confers increased lifespan independently of dietary restriction. Proc. Natl. Acad. Sci. 107, 9105–9110. 10.1073/pnas.0911539107.

21. Sanz, A., Stefanatos, R., and McIlroy, G. (2010). Production of reactive oxygen species by the mitochondrial electron transport chain in Drosophila melanogaster. J. Bioenerg. Biomembr. 42, 135–142. 10.1007/s10863-010-9281-z.

22. Chandel, N.S., McClintock, D.S., Feliciano, C.E., Wood, T.M., Melendez, J.A., Rodriguez, A.M., and Schumacker, P.T. (2000). Reactive Oxygen Species Generated at Mitochondrial Complex III Stabilize Hypoxia-inducible Factor-1α during Hypoxia. J. Biol. Chem. 275, 25130–25138. 10.1074/jbc.M001914200.

23. Movafagh, S., Crook, S., and Vo, K. (2015). Regulation of Hypoxia-Inducible Factor-1a by Reactive Oxygen Species: New Developments in an Old Debate: Regulation of Hypoxia-Inducible Factor-1a. J. Cell. Biochem. 116, 696–703. 10.1002/jcb.25074.

24. Qutub, A.A., and Popel, A.S. (2008). Reactive Oxygen Species Regulate Hypoxia-Inducible Factor 1α Differentially in Cancer and Ischemia. Mol. Cell. Biol. 28, 5106–5119. 10.1128/MCB.00060-08.

25. Huang, X., Zhao, L., and Peng, R. (2022). Hypoxia-Inducible Factor 1 and Mitochondria: An Intimate Connection. Biomolecules 13, 50. 10.3390/biom13010050.

26. Tomita, S., Ueno, M., Sakamoto, M., Kitahama, Y., Ueki, M., Maekawa, N., Sakamoto, H., Gassmann, M., Kageyama, R., Ueda, N., et al. (2003). Defective Brain Development in Mice Lacking the *Hif-1* α Gene in Neural Cells. Mol. Cell. Biol. 23, 6739–6749. 10.1128/MCB.23.19.6739-6749.2003.

27. Semenza, G.L. (2006). Regulation of physiological responses to continuous and intermittent hypoxia by hypoxia-inducible factor 1. Exp. Physiol. 91, 803–806. 10.1113/expphysiol.2006.033498.

28. Milosevic, J., Maisel, M., Wegner, F., Leuchtenberger, J., Wenger, R.H., Gerlach, M., Storch, A., and Schwarz, J. (2007). Lack of Hypoxia-Inducible Factor-1α Impairs Midbrain Neural Precursor Cells Involving Vascular Endothelial Growth Factor Signaling. J. Neurosci. 27, 412–421. 10.1523/JNEUROSCI.2482-06.2007.

29. Iacobini, C., Vitale, M., Pugliese, G., and Menini, S. (2023). The “sweet” path to cancer: focus on cellular glucose metabolism. Front. Oncol. 13, 1202093. 10.3389/fonc.2023.1202093.

30. Oswald, M.C., Brooks, P.S., Zwart, M.F., Mukherjee, A., West, R.J., Giachello, C.N., Morarach, K., Baines, R.A., Sweeney, S.T., and Landgraf, M. (2018). Reactive oxygen species regulate activity-dependent neuronal plasticity in Drosophila. eLife 7, e39393. 10.7554/eLife.39393.

31. Sobrido-Cameán, D., Oswald, M.C.W., Bailey, D.M.D., Mukherjee, A., and Landgraf, M. (2023). Activity-regulated growth of motoneurons at the neuromuscular junction is mediated by NADPH oxidases. Front. Cell. Neurosci. 16, 1106593. 10.3389/fncel.2022.1106593.

32. McDonald, A.E., and Vanlerberghe, G.C. (2004). Branched Mitochondrial Electron Transport in the Animalia: Presence of Alternative Oxidase in Several Animal Phyla. IUBMB Life 56, 333–341. 10.1080/1521-6540400000876.

33. McDonald, A.E., Vanlerberghe, G.C., and Staples, J.F. (2009). Alternative oxidase in animals: unique characteristics and taxonomic distribution. J. Exp. Biol. 212, 2627–2634. 10.1242/jeb.032151.

34. Matus-Ortega, M.G., Salmerón-Santiago, K.G., Flores-Herrera, O., Guerra-Sánchez, G., Martínez, F., Rendón, J.L., and Pardo, J.P. (2011). The alternative NADH dehydrogenase is present in mitochondria of some animal taxa. Comp. Biochem. Physiol. Part D Genomics Proteomics 6, 256–263. 10.1016/j.cbd.2011.05.002.

35. Weaver, R.J., and McDonald, A.E. (2023). Mitochondrial alternative oxidase across the tree of life: Presence, absence, and putative cases of lateral gene transfer. Biochim. Biophys. Acta BBA - Bioenerg. 1864, 149003. 10.1016/j.bbabio.2023.149003.

36. Maxwell, D.P., Wang, Y., and McIntosh, L. (1999). The alternative oxidase lowers mitochondrial reactive oxygen production in plant cells. Proc. Natl. Acad. Sci. 96, 8271– 8276. 10.1073/pnas.96.14.8271.

37. Fernandez-Ayala, D.J.M., Sanz, A., Vartiainen, S., Kemppainen, K.K., Babusiak, M., Mustalahti, E., Costa, R., Tuomela, T., Zeviani, M., Chung, J., et al. (2009). Expression of the Ciona intestinalis Alternative Oxidase (AOX) in Drosophila Complements Defects in Mitochondrial Oxidative Phosphorylation. Cell Metab. 9, 449–460. 10.1016/j.cmet.2009.03.004.

38. El-Khoury, R., Dufour, E., Rak, M., Ramanantsoa, N., Grandchamp, N., Csaba, Z., Duvillié, B., Bénit, P., Gallego, J., Gressens, P., et al. (2013). Alternative Oxidase Expression in the Mouse Enables Bypassing Cytochrome c Oxidase Blockade and Limits Mitochondrial ROS Overproduction. PLoS Genet. 9, e1003182. 10.1371/journal.pgen.1003182.

39. Pfeiffer, B.D., Jenett, A., Hammonds, A.S., Ngo, T.-T.B., Misra, S., Murphy, C., Scully, A., Carlson, J.W., Wan, K.H., Laverty, T.R., et al. (2008). Tools for neuroanatomy and neurogenetics in *Drosophila*. Proc. Natl. Acad. Sci. 105, 9715–9720. 10.1073/pnas.0803697105.

40. Liu, K., Jones, S., Minis, A., Rodriguez, J., Molina, H., and Steller, H. (2019). PI31 Is an Adaptor Protein for Proteasome Transport in Axons and Required for Synaptic Development. Dev. Cell 50, 509–524.e10. 10.1016/j.devcel.2019.06.009.

41. McClure, C.D., Hassan, A., Aughey, G.N., Butt, K., Estacio-Gómez, A., Duggal, A., Ying Sia, C., Barber, A.F., and Southall, T.D. (2022). An auxin-inducible, GAL4-compatible, gene expression system for Drosophila. eLife 11, e67598. 10.7554/eLife.67598.

42. Weidemann, A., and Johnson, R.S. (2008). Biology of HIF-1α. Cell Death Differ. 15, 621–627. 10.1038/cdd.2008.12.

43. Strowitzki, M., Cummins, E., and Taylor, C. (2019). Protein Hydroxylation by Hypoxia-Inducible Factor (HIF) Hydroxylases: Unique or Ubiquitous? Cells 8, 384. 10.3390/cells8050384.

44. Chandel, N.S., Maltepe, E., Goldwasser, E., Mathieu, C.E., Simon, M.C., and Schumacker, P.T. (1998). Mitochondrial reactive oxygen species trigger hypoxia-induced transcription. Proc. Natl. Acad. Sci. 95, 11715–11720. 10.1073/pnas.95.20.11715.

45. Brunelle, J.K., Bell, E.L., Quesada, N.M., Vercauteren, K., Tiranti, V., Zeviani, M., Scarpulla, R.C., and Chandel, N.S. (2005). Oxygen sensing requires mitochondrial ROS but not oxidative phosphorylation. Cell Metab. 1, 409–414. 10.1016/j.cmet.2005.05.002.

46. Guzy, R.D., Hoyos, B., Robin, E., Chen, H., Liu, L., Mansfield, K.D., Simon, M.C., Hammerling, U., and Schumacker, P.T. (2005). Mitochondrial complex III is required for hypoxia-induced ROS production and cellular oxygen sensing. Cell Metab. 1, 401–408. 10.1016/j.cmet.2005.05.001.

47. Fuhrmann, D.C., and Brüne, B. (2017). Mitochondrial composition and function under the control of hypoxia. Redox Biol. 12, 208–215. 10.1016/j.redox.2017.02.012.

48. Krick, N., Davies, J., Coulson, B., Sobrido-Cameán, D., Miller, M., Oswald, M.C.W., Zarin, A.A., Baines, R., and Landgraf, M. (2024). Heterogeneous responses to embryonic critical period perturbations among different components of the *Drosophila* larval locomotor circuit. Preprint, 10.1101/2024.09.14.613036 https://doi.org/10.1101/2024.09.14.613036.

49. Andersen, S.L. (2003). Trajectories of brain development: point of vulnerability or window of opportunity? Neurosci. Biobehav. Rev. 27, 3–18. 10.1016/S0149-7634(03)00005-8.

50. Cicchetti, D. (2010). Resilience under conditions of extreme stress: a multilevel perspective. World Psychiatry 9, 145–154. 10.1002/j.2051-5545.2010.tb00297.x.

51. Walton, K.D., Lieberman, D., Llinás, A., Begin, M., and Llinás, R.R. (1992). Identification of a critical period for motor development in neonatal rats. Neuroscience 51, 763–767. 10.1016/0306-4522(92)90517-6.

52. Moorman, S.J., Cordova, R., and Davies, S.A. (2002). A critical period for functional vestibular development in zebrafish. Dev. Dyn. 223, 285–291. 10.1002/dvdy.10052.

53. Avitan, L., Pujic, Z., Mölter, J., Van De Poll, M., Sun, B., Teng, H., Amor, R., Scott, E.K., and Goodhill, G.J. (2017). Spontaneous Activity in the Zebrafish Tectum Reorganizes over Development and Is Influenced by Visual Experience. Curr. Biol. 27, 2407–2419.e4. 10.1016/j.cub.2017.06.056.

54. Xie, J., Jusuf, P.R., Bui, B.V., and Goodbourn, P.T. (2019). Experience-dependent development of visual sensitivity in larval zebrafish. Sci. Rep. 9, 18931. 10.1038/s41598-019-54958-6.

55. Chin, J.S.R., Phan, T.-A.N., Albert, L.T., Keene, A.C., and Duboué, E.R. (2022). Long lasting anxiety following early life stress is dependent on glucocorticoid signaling in zebrafish. Sci. Rep. 12, 12826. 10.1038/s41598-022-16257-5.

56. Hageter, J., Starkey, J., and Horstick, E.J. (2023). Thalamic regulation of a visual critical period and motor behavior. Cell Rep. 42, 112287. 10.1016/j.celrep.2023.112287.

57. Kiral, F.R., Dutta, S.B., Linneweber, G.A., Hilgert, S., Poppa, C., Duch, C., Von Kleist, M., Hassan, B.A., and Hiesinger, P.R. (2021). Brain connectivity inversely scales with developmental temperature in Drosophila. Cell Rep. 37, 110145. 10.1016/j.celrep.2021.110145.

58. Falibene, A., Roces, F., Rössler, W., and Groh, C. (2016). Daily Thermal Fluctuations Experienced by Pupae via Rhythmic Nursing Behavior Increase Numbers of Mushroom Body Microglomeruli in the Adult Ant Brain. Front. Behav. Neurosci. 10. 10.3389/fnbeh.2016.00073.

59. Groh, C., Tautz, J., and Rössler, W. (2004). Synaptic organization in the adult honey bee brain is influenced by brood-temperature control during pupal development. Proc. Natl. Acad. Sci. 101, 4268–4273. 10.1073/pnas.0400773101.

60. Wang, G., Gordon, T.N., and Rainwater, S. (2008). Maximum voluntary temperature of insect larvae reveals differences in their thermal biology. J. Therm. Biol. 33, 380–384. 10.1016/j.jtherbio.2008.06.002.

61. Züfle, P., Batista, L.L., Brandão, S.C., D’Uva, G., Christian, D., and Martelli, C. (2023). Metabolic constraints on growth explain how developmental temperature scales synaptic connectivity relevant for behaviour. Preprint, 10.1101/2023.10.15.562389 https://doi.org/10.1101/2023.10.15.562389.

62. Crisp, S., Evers, J.F., Fiala, A., and Bate, M. (2008). The development of motor coordination in *Drosophila* embryos. Development 135, 3707–3717. 10.1242/dev.026773.

63. Crisp, S.J., Evers, J.F., and Bate, M. (2011). Endogenous Patterns of Activity Are Required for the Maturation of a Motor Network. J. Neurosci. 31, 10445–10450. 10.1523/JNEUROSCI.0346-11.2011.

64. Coulson, B., Hunter, I., Doran, S., Parkin, J., Landgraf, M., and Baines, R.A. (2022). Critical periods in Drosophila neural network development: Importance to network tuning and therapeutic potential. Front. Physiol. 13, 1073307. 10.3389/fphys.2022.1073307.

65. Graham, C., Stefanatos, R., Yek, A.E.H., Spriggs, R.V., Loh, S.H.Y., Uribe, A.H., Zhang, T., Martins, L.M., Maddocks, O.D.K., Scialo, F., et al. (2022). Mitochondrial ROS signalling requires uninterrupted electron flow and is lost during ageing in flies. GeroScience. 10.1007/s11357-022-00555-x.

66. Weidenmüller, A., Mayr, C., Kleineidam, C.J., and Roces, F. (2009). Preimaginal and Adult Experience Modulates the Thermal Response Behavior of Ants. Curr. Biol. 19, 1897– 1902. 10.1016/j.cub.2009.08.059.

67. Becher, M.A., Scharpenberg, H., and Moritz, R.F.A. (2009). Pupal developmental temperature and behavioral specialization of honeybee workers (Apis mellifera L.). J. Comp. Physiol. A 195, 673–679. 10.1007/s00359-009-0442-7.

68. Tautz, J., Maier, S., Groh, C., Rössler, W., and Brockmann, A. (2003). Behavioral performance in adult honey bees is influenced by the temperature experienced during their pupal development. Proc. Natl. Acad. Sci. 100, 7343–7347. 10.1073/pnas.1232346100.

69. Mòdol, L., Moissidis, M., Selten, M., Oozeer, F., and Marín, O. (2024). Somatostatin interneurons control the timing of developmental desynchronization in cortical networks. Neuron 112, 2015–2030.e5. 10.1016/j.neuron.2024.03.014.

70. Tien, N.-W., and Kerschensteiner, D. (2018). Homeostatic plasticity in neural development. Neural Develop. 13, 9. 10.1186/s13064-018-0105-x.

71. Chen, L., Li, X., Tjia, M., and Thapliyal, S. (2022). Homeostatic plasticity and excitation-inhibition balance: The good, the bad, and the ugly. Curr. Opin. Neurobiol. 75, 102553. 10.1016/j.conb.2022.102553.

72. Swann, J.W., and Rho, J.M. (2014). How Is Homeostatic Plasticity Important in Epilepsy? In Issues in Clinical Epileptology: A View from the Bench Advances in Experimental Medicine and Biology., H. E. Scharfman and P. S. Buckmaster, eds. (Springer Netherlands), pp. 123–131. 10.1007/978-94-017-8914-1_10.

73. Davis, G.W., and Bezprozvanny, I. (2001). Maintaining the Stability of Neural Function: A Homeostatic Hypothesis. Annu. Rev. Physiol. 63, 847–869. 10.1146/annurev.physiol.63.1.847.

74. O’Donovan, M.J., Chub, N., and Wenner, P. (1998). Mechanisms of spontaneous activity in developing spinal networks. J. Neurobiol. 37, 131–145. 10.1002/(SICI)1097-4695(199810)37:1<131::AID-NEU10>3.0.CO;2-H.

75. Wong, W.T., Sanes, J.R., and Wong, R.O.L. (1998). Developmentally Regulated Spontaneous Activity in the Embryonic Chick Retina. J. Neurosci. 18, 8839–8852. 10.1523/JNEUROSCI.18-21-08839.1998.

76. Dhawan, S., Myers, P., Bailey, D.M.D., Ostrovsky, A.D., Evers, J.F., and Landgraf, M. (2021). Reactive Oxygen Species Mediate Activity-Regulated Dendritic Plasticity Through NADPH Oxidase and Aquaporin Regulation. Front. Cell. Neurosci. 15, 641802. 10.3389/fncel.2021.641802.

77. Robb, E.L., Hall, A.R., Prime, T.A., Eaton, S., Szibor, M., Viscomi, C., James, A.M., and Murphy, M.P. (2018). Control of mitochondrial superoxide production by reverse electron transport at complex I. J. Biol. Chem. 293, 9869–9879. 10.1074/jbc.RA118.003647.

78. Truszkowski, T.L.S., and Aizenman, C.D. (2015). Neurobiology: Setting the Set Point for Neural Homeostasis. Curr. Biol. 25, R1132–R1133. 10.1016/j.cub.2015.10.021.

79. Styr, B., Gonen, N., Zarhin, D., Ruggiero, A., Atsmon, R., Gazit, N., Braun, G., Frere, S., Vertkin, I., Shapira, I., et al. (2019). Mitochondrial Regulation of the Hippocampal Firing Rate Set Point and Seizure Susceptibility. Neuron 102, 1009–1024.e8. 10.1016/j.neuron.2019.03.045.

80. Vander Heiden, M.G., Cantley, L.C., and Thompson, C.B. (2009). Understanding the Warburg Effect: The Metabolic Requirements of Cell Proliferation. Science 324, 1029–1033. 10.1126/science.1160809.

81. Wang, C.-W., Purkayastha, A., Jones, K.T., Thaker, S.K., and Banerjee, U. (2016). In vivo genetic dissection of tumor growth and the Warburg effect. eLife 5, e18126. 10.7554/eLife.18126.

82. Elzakra, N., and Kim, Y. (2021). HIF-1α Metabolic Pathways in Human Cancer. In Cancer Metabolomics Advances in Experimental Medicine and Biology., S. Hu, ed. (Springer International Publishing), pp. 243–260. 10.1007/978-3-030-51652-9_17.

83. Dunwoodie, S.L. (2009). The Role of Hypoxia in Development of the Mammalian Embryo. Dev. Cell 17, 755–773. 10.1016/j.devcel.2009.11.008.

84. Bohuslavova, R., Cerychova, R., Papousek, F., Olejnickova, V., Bartos, M., Görlach, A., Kolar, F., Sedmera, D., Semenza, G.L., and Pavlinkova, G. (2019). HIF-1α is required for development of the sympathetic nervous system. Proc. Natl. Acad. Sci. 116, 13414– 13423. 10.1073/pnas.1903510116.

85. Kallio, P.J. (1998). Signal transduction in hypoxic cells: inducible nuclear translocation and recruitment of the CBP/p300 coactivator by the hypoxia-inducible factor-1alpha. EMBO J. 17, 6573–6586. 10.1093/emboj/17.22.6573.

86. Perez-Perri, J.I., Dengler, V.L., Audetat, K.A., Pandey, A., Bonner, E.A., Urh, M., Mendez, J., Daniels, D.L., Wappner, P., Galbraith, M.D., et al. (2016). The TIP60 Complex Is a Conserved Coactivator of HIF1A. Cell Rep. 16, 37–47. 10.1016/j.celrep.2016.05.082.

87. Luo, W., and Wang, Y. (2018). Epigenetic regulators: multifunctional proteins modulating hypoxia-inducible factor-α protein stability and activity. Cell. Mol. Life Sci. 75, 1043–1056. 10.1007/s00018-017-2684-9.

88. Yfantis, A., Mylonis, I., Chachami, G., Nikolaidis, M., Amoutzias, G.D., Paraskeva, E., and Simos, G. (2023). Transcriptional Response to Hypoxia: The Role of HIF-1-Associated Co-Regulators. Cells 12, 798. 10.3390/cells12050798.

89. Sulkowski, M.J., Han, T.H., Ott, C., Wang, Q., Verheyen, E.M., Lippincott-Schwartz, J., and Serpe, M. (2016). A Novel, Noncanonical BMP Pathway Modulates Synapse Maturation at the Drosophila Neuromuscular Junction. PLOS Genet. 12, e1005810. 10.1371/journal.pgen.1005810.

90. Bataillé, L., Delon, I., Da Ponte, J. P., Brown, N. H., & Jagla, K. (2010). Downstream of identity genes: muscle-type-specific regulation of the fusion process. Developmental cell, 19(2), 317–328. 10.1016/j.devcel.2010.07.008

91. Fujioka, M., Lear, B. C., Landgraf, M., Yusibova, G. L., Zhou, J., Riley, K. M., et al. (2003). Even-skipped, acting as a repressor, regulates axonal projections in Drosophila. Development 130, 5385–5400. doi: 10.1242/dev.00770

